# Revealing the Drivers Underlying Distinct Evolutionary Trajectories in Lung Adenocarcinoma

**DOI:** 10.64898/2025.12.19.695410

**Authors:** Christopher Wirth, Tongwu Zhang, Marcos Díaz-Gay, Christopher D. Steele, Phuc H. Hoang, Yang Yang, Azhar Khandekar, Wei Zhao, Jian Sang, Charles Leduc, Marina K. Baine, William D. Travis, Lynette M. Sholl, Philippe Joubert, Robert Homer, Soo-Ryum Yang, Thi-Van-Trinh Tran, John P. McElderry, Caleb Hartman, Mona Miraftab, Olivia W. Lee, Kristine M. Jones, Bin Zhu, Jacobo Martínez Santamaría, Matthew B. Schabath, Sai S. Yendamuri, Marta Manczuk, Jolanta Lissowska, Beata Świątkowska, Anush Mukeria, Oxana Shangina, David Zaridze, Ivana Holcatova, Vladimir Janout, Dana Mates, Simona Ognjanovic, Milan Savic, Milica Kontic, Yohan Bossé, Bonnie E. Gould Rothberg, David C. Christiani, Valerie Gaborieau, Paul Brennan, Geoffrey Liu, Paul Hofman, Maria Pik Wong, Kin Chung Leung, Chih-Yi Chen, I-Shou Chang, Chao Agnes Hsiung, Angela C. Pesatori, Dario Consonni, Nathaniel Rothman, Qing Lan, Martin A. Nowak, Stephen J. Chanock, Jianxin Shi, Lixing Yang, Ludmil B. Alexandrov, David C. Wedge, Maria Teresa Landi

## Abstract

Elucidating the evolution of cancers allows us to understand their key events, and the order in which they occur. To chart and interpret these evolutionary trajectories, we leverage whole-genome sequencing of lung tumours, including those from the largest cohort to date of lung cancers in subjects who have never smoked. Through ordering frequent genomic alterations, we discover three distinct evolutionary paths taken by lung adenocarcinomas; two dominated by tumours from people who have never smoked (NS-LUAD), and one followed by the vast majority of those who have smoked (S-LUAD). However, one in six NS-LUAD follow the smoking-dominant trajectory. These tumours, surprisingly, have fewer somatic alterations than the other NS-LUAD, and have shorter latency. They are strongly enriched for *KRAS* mutations. Our results suggest that gaining *KRAS* mutations allows these tumours to evolve more rapidly, acquiring a set of smoking-associated key alterations, with less need for genomic instability to progress. These tumours are three times more frequent in subjects of European vs. East Asian ancestry. These findings could shape clinical management strategies for lung adenocarcinoma patients, particularly for tumours driven by smoking-like evolutionary trajectories.

## Main

Understanding the biological processes of lung cancer in people who have never smoked (LCINS) is a matter of growing importance in the field of cancer research. The proportion of lung cancer patients who have never smoked is increasing^1,2^, likely due to a general trend towards non-smoking at the population level, particularly in younger people^3^. There is therefore a pressing and increasing need to understand the development and evolution of the disease in people who have never smoked.

Charting the evolutionary trajectories of cancers allows us to understand the key events, and the order in which they occur, throughout tumour development. A portion of the LCINS cohort included in this paper (n=232) has previously been analysed^4^, estimating the order of events in subsets of samples with similar genomic profiles at the time of sampling. This identified mutations to driver genes in the RTK-RAS pathway, including *TP53*, *EGFR*, *KRAS*, and *RMB10* as common early driver events, as well as early copy number losses that were specific to one subset. More recently, de novo discovery of subsets through the clustering of event orderings has been utilised as a method for uncovering distinct evolutionary subtypes of prostate cancer^5^. Explicitly taking into account the order of events in each tumour when grouping samples yields important and hitherto unforeseen insights. Whereas clustering samples according to their static genomic profiles at the time of sampling is likely to group together samples with similar mutation burdens and driver gene type, evolutionary subtyping is likely to group together tumours at different stages of their evolution, enabling the inference of the key events that lead to progression from a less advanced to a more advanced tumour.

In this study we conducted an in-depth investigation of the evolutionary trajectories of LCINS, leveraging the expanded Sherlock-*Lung* dataset^6^. To minimise histology-related bias in our *de novo* stratification, we focused on lung adenocarcinomas (LUAD). The full Sherlock-*Lung* whole-genome sequencing sample set (n=1217) was filtered to LUAD samples with at least 10 reads per tumour chromosome copy^7^ and a minimum tumour purity of 30%. This yielded a cohort of 550 LUAD tumours with sufficient power to reliably identify subclones with cancer cell fraction (CCF) over 40%^8^. Of these, 407 were self-reported as coming from patients who had never smoked (hereafter referred to as NS-LUAD) and 143 from patients who had smoked (hereafter referred to as S-LUAD); 369 were female and 181 male; 339 were assigned European genetic ancestry and 190 East Asian genetic ancestry, with a further 21 assigned other genetic ancestries (**Methods**). Further demographic information including the intersections between these factors is shown in **Table S1**.

We used the Plackett-Luce ordering model that we previously developed^5^ (**Methods**) to perform *de novo* discovery of evolutionary trajectories, identifying three evolutionary paths with distinct genomic and clinical associations. For each trajectory, we calculated average copy number profiles across the corresponding subsets of tumours and collated subset-wide alteration metrics, allowing us to investigate and compare the genomic states that had been reached following the accumulation of somatic events. We further examined preferred clonal aberrations to pinpoint likely disease-initiating events, and the key drivers associated with each trajectory. Finally, we investigated the impact of early driver mutations on genomic instability.

These analyses allowed us to construct a full picture of the varying modes of evolution in LUAD. Importantly, they revealed specific genomic events that can drive NS-LUAD to evolve in a smoking-resemblant manner.

## Results

### *De novo* discovery identifies three evolutionary trajectories in LUAD

Frequently occurring somatic copy number alterations (SCNAs) and single nucleotide variants (SNVs) were identified and ordered, and subsets of samples with related evolutionary trajectories were discovered using a Plackett-Luce model. The steps of the model are illustrated in **Fig. 1a**. This *de novo* ordering analysis identified three subsets of LUAD (**Fig. 1b, Fig. S1, Methods**, **Table S2**). The first two subsets are predominantly composed of NS-LUAD (96% and 91% respectively) and are therefore referred to as NS-LUAD Dominant (NSD) trajectories. In contrast, the third subset includes the majority (84%) of S-LUAD and is thus designated as the S-LUAD Dominant trajectory (SD). Notably, this subset is less exclusive than the NSD subsets; 18% of NS-LUAD tumours are classified within the SD trajectory, accounting for over one third of its total composition. One NS-LUAD sample could not be categorised as belonging to any subset, as it did not contain any of the commonly occurring SCNAs, or mutations in driver genes. This sample was therefore excluded from downstream analyses.

**Fig. 1:**
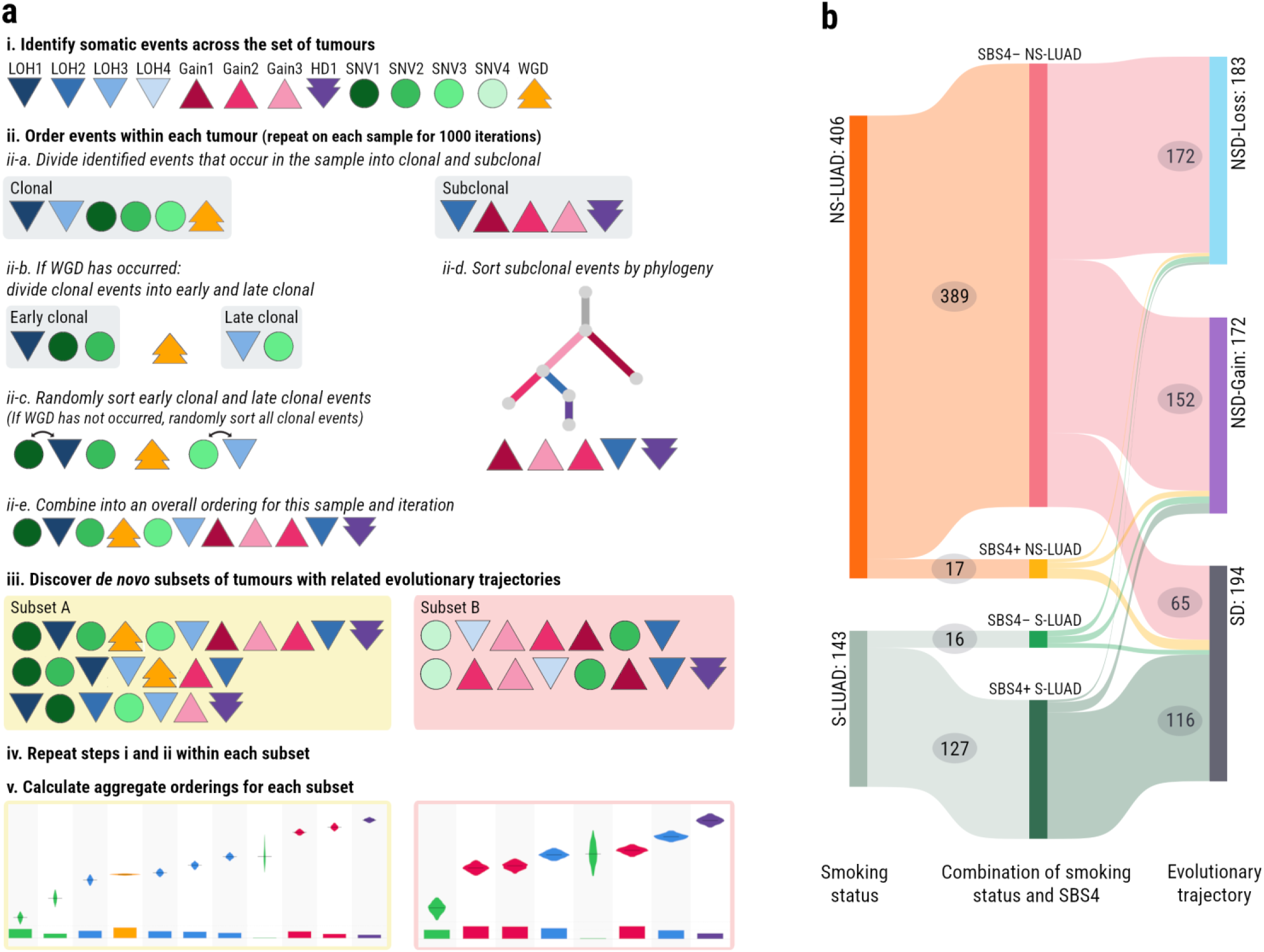
The Plackett-Luce model and *de novo* subsetting. **a)** An illustration of the Plackett-Luce model for *de novo* discovery of subsets and ordering of events. SCNAs occurring more frequently than expected by chance are identified using binomial probability. SNVs in a pre-specified list of driver genes are included in the model if they occur in >= 3% of samples. In this illustration, LOH events are identified by downward pointing triangles in different shades of blue, copy number gains are identified by upward pointing triangles in different shades of blue, homozygous deletions are identified by downward arrows in purple, and WGD is identified by an upward arrow in orange. For each tumour and each of 1000 iterations: Events are divided into clonal & subclonal, clonal events in WGD samples are divided into early & late clonal and events are randomly ordered within these sets, subclonal events are ordered by phylogeny, and the events are then joined into a complete order. *De novo* subsets of tumours with similar sets and orders of events are discovered. The earlier steps of the model are then carried out again on each subset of tumours before an aggregate ordering is calculated and plotted for each *de novo* trajectory. **b)** A Sankey-style diagram of the *de novo* discovered LUAD subsets in the context of smoking status and the presence or absence of SBS4 mutations. 406 NS-LUAD tumours are divided into 389 SBS4- and 17 SBS4+ tumours. 143 S-LUAD tumours are divided into 16 SBS4- tumours and 127 SBS4+ tumours. Of the SBS4- NS-LUAD samples, 172 are assigned to NSD-Loss, 152 to NSD-Gain, and 65 to SD. 116 SBS4+ tumours are assigned to SD. All other paths contain fewer than 10 samples and these numbers are not shown on the diagram.

The three subsets follow distinct trajectories of events (**Fig. 2**). The first NSD subset features whole-genome duplication (WGD) in 87% of its samples (**Fig. 2a**). WGD generally occurs early in the evolution of these tumours, with only six key events - three driver genes *(RBM10, EGFR, TP53)* bearing SNVs and three copy number losses (including 17p13.3-11.2/*TP53*) - generally preceding it in this trajectory. The trajectory is further characterised by a striking dichotomy in copy number activity: losses dominate the early activity, followed by gains occurring later and at lower prevalence. We therefore refer to this as the NSD-Loss trajectory. In contrast, the second NSD subset is predominantly composed of non-WGD samples (69%) (**Fig. 2b**) and where WGD is present, it generally occurs later than in NSD-Loss tumours. The ploidy of these tumours is more likely to increase via early, frequent copy number gains than through WGD. We therefore refer to this as the NSD-Gain trajectory. The order of copy number events in the SD subset (**Fig. 2c**) shows a similar pattern to the NSD-Loss trajectory, where all loss of heterozygosity (LOH) events precede all gains within the ordering. WGD is also an early event in the SD trajectory; although it is preceded by many more driver genes bearing SNVs, again only three copy number losses (also including 17p13.3-11.2/*TP53*) generally occur prior to WGD.

**Fig. 2:**
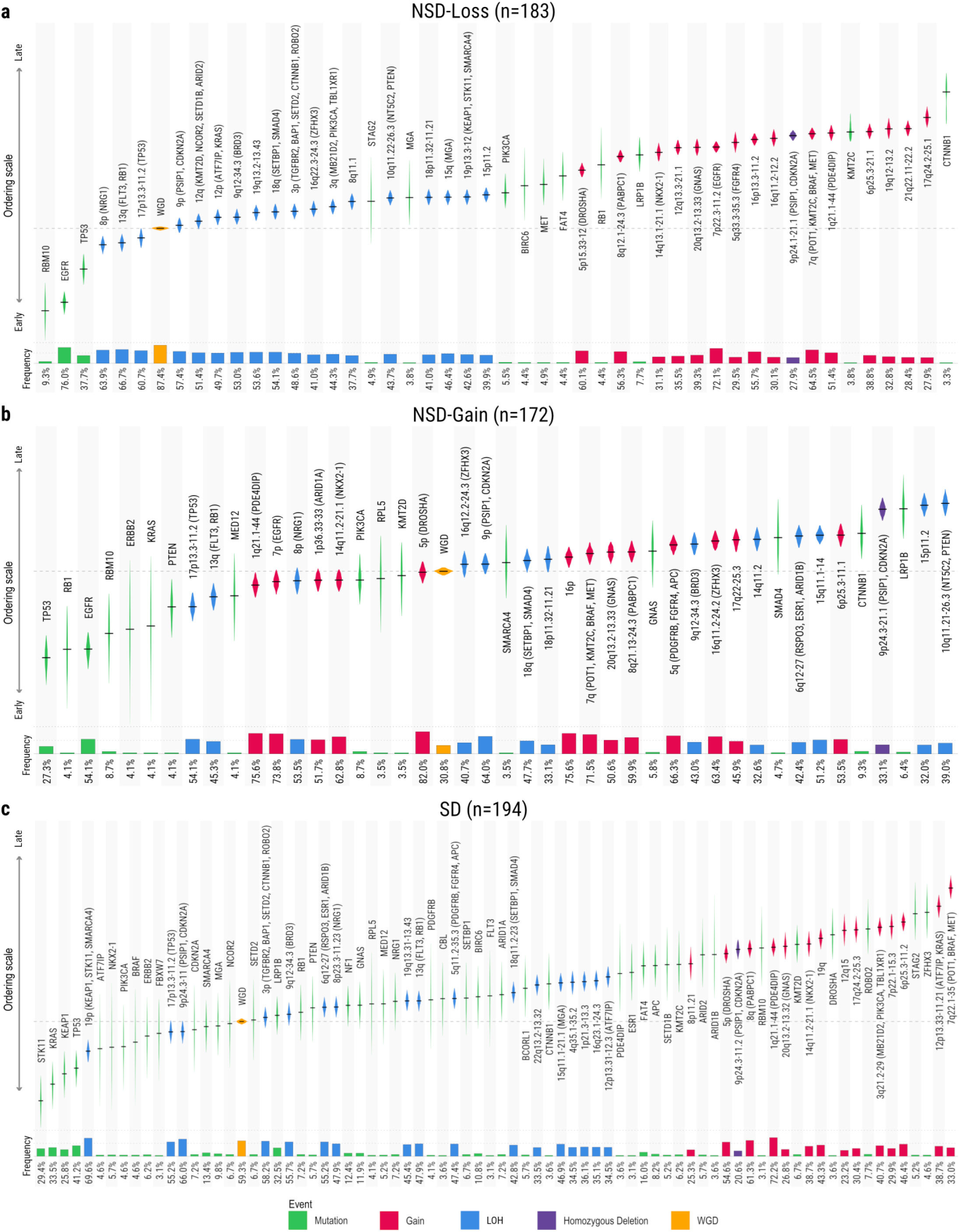
***De novo* evolutionary orderings. a)** The *NSD-Loss* ordering (n=183). **b)** The *NSD-Gain* ordering (n=172). **c)** The *SD* ordering (n=194). In all orderings, events are ordered by an aggregate ordering index, from earliest at the bottom/left to latest at the top/right. The extent of each violin plot shows the 95% confidence interval of the event’s aggregate ordering, across 1000 iterations of the Plackett-Luce algorithm. Nonsynonymous SNVs in driver genes are labelled with the name of the gene, and displayed in green. SCNAs are labelled with the altered cytobands, with any driver genes contained within the segments included in brackets. Losses are displayed in blue, gains in red, and homozygous deletions in purple. Whole genome duplication is labelled “WGD” and displayed in orange. The size of the bars beneath the violin plots represent the proportion of samples in the subset that carry the event.

All subsets acquire driver mutations as the earliest events in the evolutionary ordering, preceding any SCNAs. In both NSD trajectories, the most common early mutations occur in *EGFR* and *TP53*, with notable less frequent mutations observed in *ERBB2*, *KRAS*, and *RBM10*. In contrast, the SD trajectory shows a distinct pattern: *EGFR* mutations are very rare, occurring in just 2% of samples, yet *TP53* mutations remain a frequent early event. Early mutations in *STK11*, *KRAS*, and *KEAP1* also occur in a substantial portion of SD samples. Rarer mutations occur throughout tumour development of all trajectories, but substantially more of these mutations are observed in the SD trajectory (45 driver genes contained SNVs in >= 3% of SD tumours vs 13 in NSD-Loss and 16 in NSD-Gain, FDR=3.7×10^-9^, **Methods**). These observations are in line with our expectations given the relative enrichment of NS-LUAD and S-LUAD across trajectories and the increased tumour mutation burden (TMB) associated with tobacco smoking^9–11^. The substantial portion of NS-LUAD tumours assigned to the SD trajectory, featuring these known smoking-related alterations, is of great interest.

The loss on 17p (*TP53*) is the only SCNA consistently ordered before WGD across all trajectories. This marks an important functional difference, as LOH (prior to WGD) results in complete loss of one allele, whereas a post-WGD loss would merely cause a reduction in the number of copies. Further, 96% of *TP53* mutant tumours also feature the 17p13.3-11.2 loss, placing this among the most frequently co-occurring pairs of events in our data (**Fig. S2**, **Table S3**). The consistently early ordering of both mutation and loss of *TP53* highlights its importance in tumour initiation regardless of the trajectory a tumour will follow. 9p (*CDKN2A, PSIP1*) also undergoes LOH in most samples of all subsets, and this SCNA was relatively early in all trajectories - being ordered prior to WGD in the SD trajectory and shortly after WGD in both NSD trajectories. Beyond these commonalities, other regions targeted by SCNAs show more marked differences between trajectories. The losses of 8p (*NRG1*) and 13q (*RB1, FLT3*) are the other copy number losses with pre-WGD orderings in both NSD trajectories, yet they occur later in the SD trajectory. Conversely, the loss of 19p (*STK11, KEAP1, SMARCA4*) is the earliest ordered SCNA in the SD trajectory, but one of the latest copy number losses in the NSD-Loss trajectory. This loss does not occur in more samples than expected by chance among NSD-Gain tumours according to binomial probability. Therefore, it is not included in the NSD-Gain trajectory at all (**Methods**). Similarly, the loss of 3p (*TGFBR1, BAP1, SETD2, ROBO2, CTNNB1*) is the fourth earliest LOH to appear in the aggregate ordering of the SD trajectory but appears slightly later (tenth) in the NSD-Loss trajectory, and not at all in the NSD-Gain trajectory, due to low prevalence.

Copy number gains of 1q21.1-44 (*PDE4DIP*) and 7p (*EGFR)* are the 3rd and 4th SCNAs overall, respectively, in the NSD-Gain trajectory. Both have an aggregate ordering prior to WGD. Conversely, the gain of *PDE4DIP* is ordered much later in the NSD-Loss and SD trajectories, while the gain of *EGFR* is ordered late in the NSD-Loss trajectory and is not included in the SD trajectory at all due to low prevalence in this subset.

Smoking status is the demographic metric that shows the strongest association with the evolutionary trajectories (**Table S1**, **Fig. S3a-b**). However, if we consider only NS-LUAD and compare our two largest ancestry cohorts, we find that NS-LUAD in subjects with European ancestry is almost three times as likely to evolve on the SD trajectory compared to those with East Asian ancestry (26.1% vs 9.1%, FDR=3.2×10^-5^, **Fig. S3c**). This suggests that, in addition to their smoking history, a patient’s genetic background may play a significant role in determining the evolutionary route that their tumour will take. This is likely related to the fact that *KRAS* mutations are more prevalent in NS-LUAD in European subjects than those from East Asia, whereas *EGFR* mutations are more common^12,13^. Conversely, neither sex nor passive smoking display any significant influence over evolutionary trajectories (FDR > 0.05, **Fig. S3e-g**). Among NS-LUAD tumours, we identified a few cases showing the tobacco smoking signature SBS4^14,15^. These are strongly associated with the SD trajectory (risk ratio=3.17, FDR=0.002, **Fig. S3d**). However, 65 of the 74 (88%) NS-LUAD on the SD trajectory are SBS4-negative (**Fig. 1b**). Therefore in the remainder of these analyses, where we stratify by smoking status, we look specifically at SBS4-positive S-LUAD and SBS4-negative NS-LUAD. This allows us to investigate what, in the absence of exposures resulting in SBS4^6^, prompts NS-LUAD to evolve in a smoking-like manner.

### *KRAS* and *STK11* alterations push NS-LUAD tumours to the smoking-like trajectory

Many genomic events show significant differentiation between the combined NSD trajectories and the SD trajectory (**Fig. 3a**, **Table S4**). Interestingly, only a small subset of these events appear to play a significant role in redirecting NS-LUAD tumours towards the SD trajectory (**Fig. 3b**, **Table S5**).

**Fig. 3:**
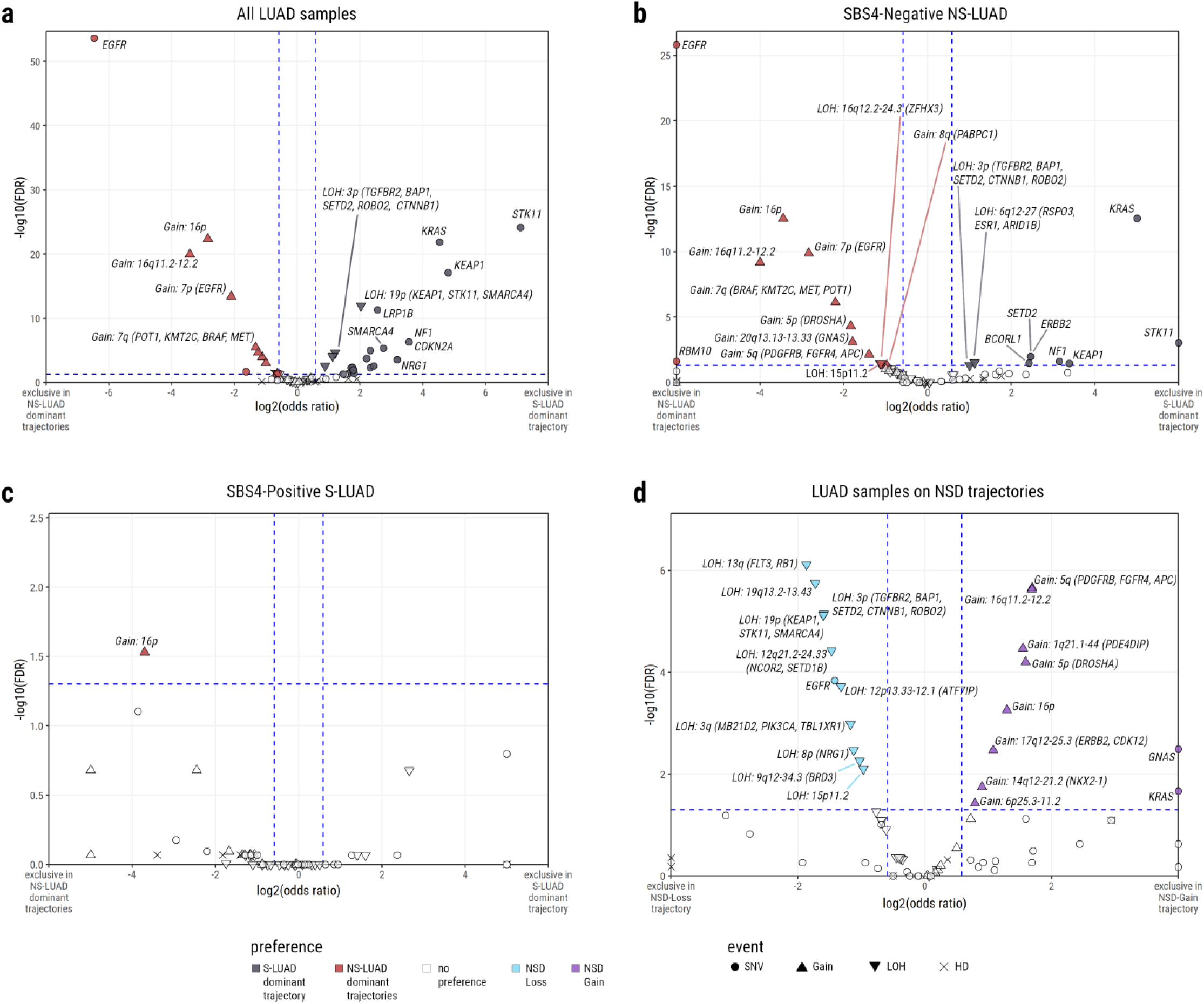
Events showing a significant preference between trajectories. Volcano plots show the log2(odds ratio) and - log10(FDR) of event occurrence comparing: **a)** All LUAD between the *SD* ordering and the *NSD-Loss* & *NSD-Gain* orderings combined. **b)** NS-LUAD between the *SD* ordering and the *NSD-Loss* & *NSD-Gain* orderings combined. **c)** S-LUAD between the *SD* ordering and the *NSD-Loss* & *NSD-Gain* orderings combined. **d)** All LUAD between the *NSD-Loss* ordering and the *NSD-Gain* ordering. Events are coloured if odds ratio > 3/2 or < 2/3, and FDR < 0.05. Blue dashed lines represent these thresholds. Events are coloured according to their positive associations as follows: SD trajectory (vs NSD trajectories combined) - black, NSD trajectories combined (vs SD trajectory) - red, NSD-Loss (vs NSD-Gain) - blue, NSD-Gain (vs NSD-Loss) - purple, no association - white. SNVs are represented by circles, copy number gains by upward pointed triangles, losses by downward pointed arrows, and HDs by crosses. In (a), due to the number of events meeting the positive association criteria, events are only labelled if odds ratio > 8 or < 1/8, or if FDR < 1^-5^. In (b)-(d), all events meeting the positive association criteria are labelled.

*KRAS* and *STK11* have been observed to be more frequently mutated in S-LUAD than NS-LUAD^16–20^, with *KRAS* mutations known to activate the MAPK pathway^16–18,20–25^. Among NS-LUAD on the SD trajectory, *KRAS* mutations are the most prominent, occurring in 22/65 tumours (34%) and representing the strongest association with the SD trajectory in this group (**Fig. 3b**, **Table S5**, OR for enrichment in SD = 32.1, FDR = 2.8×10^-13^). *STK11* mutations are rare in NS-LUAD. Where *STK11* mutations are observed within NS-LUAD, they are found exclusively in the SD trajectory (FDR = 0.0009). SNVs in *KEAP1*, *NF1*, *ERBB2*, *SETD2*, and *BCORL1* are also marginally associated with the SD trajectory in NS-LUAD (ORs for enrichment in SD in range 5.4 to 10.5, FDRs in range 0.010 to 0.034). Two LOH events also show significant associations with SD trajectory NS-LUAD tumours: loss of 6q (encompassing *ARID1B*, *ESR1*, and *RSPO3;* OR for enrichment in SD = 2.2, FDR = 0.030)—and loss of 3p (harbouring four tumour suppressors^26–29^: *SETD2*, *TGFBR2*, *BAP1*, and *ROBO2;* OR for enrichment in SD = 2.0, FDR = 0.049). Notably, both losses have been previously reported as more common in S-LUAD than NS-LUAD^30,31^.

*EGFR* is mutated more frequently in NS-LUAD than S-LUAD^16,17,20^. In accordance with this, *EGFR* mutations are the events most significantly associated with NSD trajectories across all LUAD samples (**Fig. 3a**, OR for enrichment in NSD = 88.9, FDR = 2.2×10^-54^), and found exclusively in NSD tumours when considering only NS-LUAD (**Fig. 3b**, FDR = 1.5×10^-26^). Copy number gain of *EGFR* on chromosome 7p is also strongly associated with NSD trajectories both among all LUAD (**Fig. 3a**, OR for enrichment in NSD = 4.3, FDR = 3.4×10^-14^) and among NS-LUAD (**Fig. 3b**, OR for enrichment in NSD = 7.8, FDR = 1.3×10^-10^). *EGFR* is upstream of *KRAS* on the MAPK pathway^17,20,21,25,32^ and is also influential in the activation of numerous other pathways including the PI3K and STAT pathways^25,32,33^. In addition to *EGFR*, a copy number gain including two further growth factor receptor genes—*PDGFRB* and *FGFR4* on 5q—is significantly associated with the NSD trajectories in all LUAD (**Fig. 3a**, OR for enrichment in NSD = 2.4, FDR = 2.2×10^-5^) and among NS-LUAD (**Fig. 3b**, OR for enrichment in NSD = 2.6, FDR = 0.007). This gain commonly co-occurs with the gain of *EGFR* on 7p (**Fig. S2**, **Table S3**), and generally occurs later in the trajectory (**Fig. 2**). However, 46 of 246 (19%) tumours with this 5q gain are absent of either an *EGFR* mutation or copy number gain. *FGFR4* is a known oncogene^34^ which has been shown to work in tandem with *EGFR* in LUAD^35^ and whose enhanced signalling has been shown to drive progression of squamous cell lung carcinoma^36^. Both *FGFR4* and *PDGFRB* are involved in the MAPK and PI3K pathways^25^ and have been associated with enhanced proliferation^37–41^. These findings suggest that NSD tumours may activate the MAPK pathway via distinct mechanisms from SD tumours, or may activate alternative signalling pathways altogether. Importantly, both *PDGFRB* and *FGFR4* have been investigated as potential therapeutic targets for cancer treatments^41–47^, providing possible alternative avenues for targeted treatment in patients with these alterations.

Copy number gains on both the p and q arms of chromosome 16 are associated with NSD trajectories across all LUAD (**Fig. 3a**, OR for enrichment in NSD = 7.2 and 10.7, FDR = 4.1×10^-23^ and 1.1×10^-20^, respectively) and among NS-LUAD (**Fig. 3b**, OR for enrichment in NSD = 10.9 and 16.1, FDR = 2.8×10^-13^ and 6.7×10^-10^, respectively). These large regions are frequently gained together (**Fig. S2**, **Table S3**) and may have several targets, but these remain uncertain. One such potential target is *FUS*, on 16p11.2, which it has been proposed may act as an oncogene when gained in LUAD^48^. Gains on 7q (harbouring *MET*, *BRAF*, *KMT2C*, and *POT1*), 20q (*GNAS*), and 5p (*DROSHA*) are all also associated with the NSD trajectories both across all LUAD, and when considering only NS-LUAD. Additionally, two events—a gain of 14q (*NKX2-1*) and a homozygous deletion on 9p (*CDKN2A*, *PSIP1*)—are significantly associated with NSD trajectories among all LUAD, but not specifically among NS-LUAD. Conversely, three events—losses on 16q (*ZFHX3*) and 15p, and a gain on 8q (*PABPC1*)—do not show any significant associations when considering all LUAD, but are associated with the NSD trajectories among NS-LUAD.

*FAT4* mutations are observed in 21% of S-LUAD tumours but only 3% of NS-LUAD tumours. They are significantly associated with the SD trajectory when considering all LUAD (**Table S4**), but not when restricting analysis to NS-LUAD (**Table S5**). *FAT4* has been shown to be involved in the regulation of the MAPK pathway in LUAD^49^, providing further evidence of differing mechanisms for affecting this pathway between the NSD and SD trajectories. Additionally, these findings imply that some smoking-associated mechanisms are not frequently observed in NS-LUAD, even when these tumours are evolving in a more smoking-like manner.

Other events that are significantly associated with the SD trajectory across all LUAD, but not among NS-LUAD, include mutation of *SMARCA4*, and the loss of this gene along with *STK11* and *KEAP1* on 19p. These alterations are well-documented as more prevalent in S-LUAD^50–53^ - a trend mirrored in our data (**Fig. S4**, **Table S8**). However, they do not appear to play significant roles in shifting NS-LUAD tumours toward a smoking-like trajectory.

Due to limited sample size - only 11 out of 127 S-LUAD tumours follow NSD trajectories - there is constrained statistical power to identify events that differentiate the NSD versus SD trajectories among S-LUAD. One event displays significant association: copy number gain of 16p (*FUS*) is significantly associated with the NSD trajectories among S-LUAD (**Fig. 3c**, **Table S6**, OR for enrichment in NSD = 13.0, FDR = 0.029). Additionally, *STK11* mutations were again found exclusively in SD tumours among S-LUAD, mirroring the pattern observed in NS-LUAD.

It is important to note that two distinct NSD trajectories have been identified in this study. When the NSD-Loss and NSD-Gain trajectories are compared, *EGFR* mutations are more strongly associated with NSD-Loss. Conversely, among the NSD trajectories, *KRAS* mutations are exclusive to NSD-Gain. All other events associated with NSD-Loss are copy number losses, whereas 8 of the 10 events significantly associated with NSD-Gain are copy number gains (**Fig. 3d**, **Table S7**). This reinforces the earlier description of the trajectories, as NSD-Loss is characterised by frequent WGD combined with copy number losses, whereas individual copy number gain events are observed at higher prevalence in NSD-Gain (**Fig. 2**). Many of the copy number events associated with NSD-Gain compared to NSD-Loss are those which were associated with the NSD trajectories compared to the SD trajectory; here we observe the gains on 5q (*PDGFRB*, *FGFR4*), 5p (*DROSHA*), 16p (*FUS*), and 16q. Meanwhile, some of the events associated with NSD-Loss are those which, overall, are observed more frequently in the SD trajectory than NSD trajectories. These include the losses of 3p tumour suppressors (*ROBO2*, *SETD2*, *BAP1*, *TGFBR2*) and 19p (*STK11*, *KEAP1*, *SMARCA4*). This implies that these events, which have previously been reported as less common in NS-LUAD than S-LUAD^30,31,50^, are in fact specifically rare in one subtype of NS-LUAD: the NSD-Gain trajectory.

### Copy number states are indicative of key driver events

While the trajectory analysis provides detailed event-based insights, we can also consider the overall states that each tumour had reached at the time of sampling. **Fig. 4a** shows the mean copy number state, calculated across all tumours on each trajectory. Many of the regions showing the greatest differences echo those identified in our event-based analysis. The total copy number across 7p is approximately one chromosome copy higher on average in both NSD trajectories than the SD trajectory. This difference appears to be due to the large number of NSD tumours with copy number gains of this region, encompassing *EGFR*. 16p displays a similar pattern of substantially greater total and major copy number in NSD tumours. Chromosome 5q shows striking divergence in major, minor, and total copy number states between NSD trajectories and the SD trajectory. This is the result of two factors: the more frequent gain of the region featuring *PDGFRB* and *FGFR4* on this arm in NSD trajectory tumours, and the more frequent loss of *APC* in the region in SD tumours. The average minor copy number on 19p (*STK11*, *SMARCA4*, *KEAP1*) is significantly lower in SD samples than NSD—especially in comparison with NSD-Gain. This echoes the event-based finding that this LOH is more common in SD than NSD trajectories and more common in NSD-Loss than NSD-Gain. Interestingly, the loss of tumour suppressors on 3p results in almost identical minor copy number between the NSD-Loss and SD trajectories, with both being significantly lower than NSD-Gain. This reiterates our finding from the previous section; the previously reported rarity of this event in NS-LUAD^30,31^ is particular to the NSD-Gain trajectory.

**Fig. 4:**
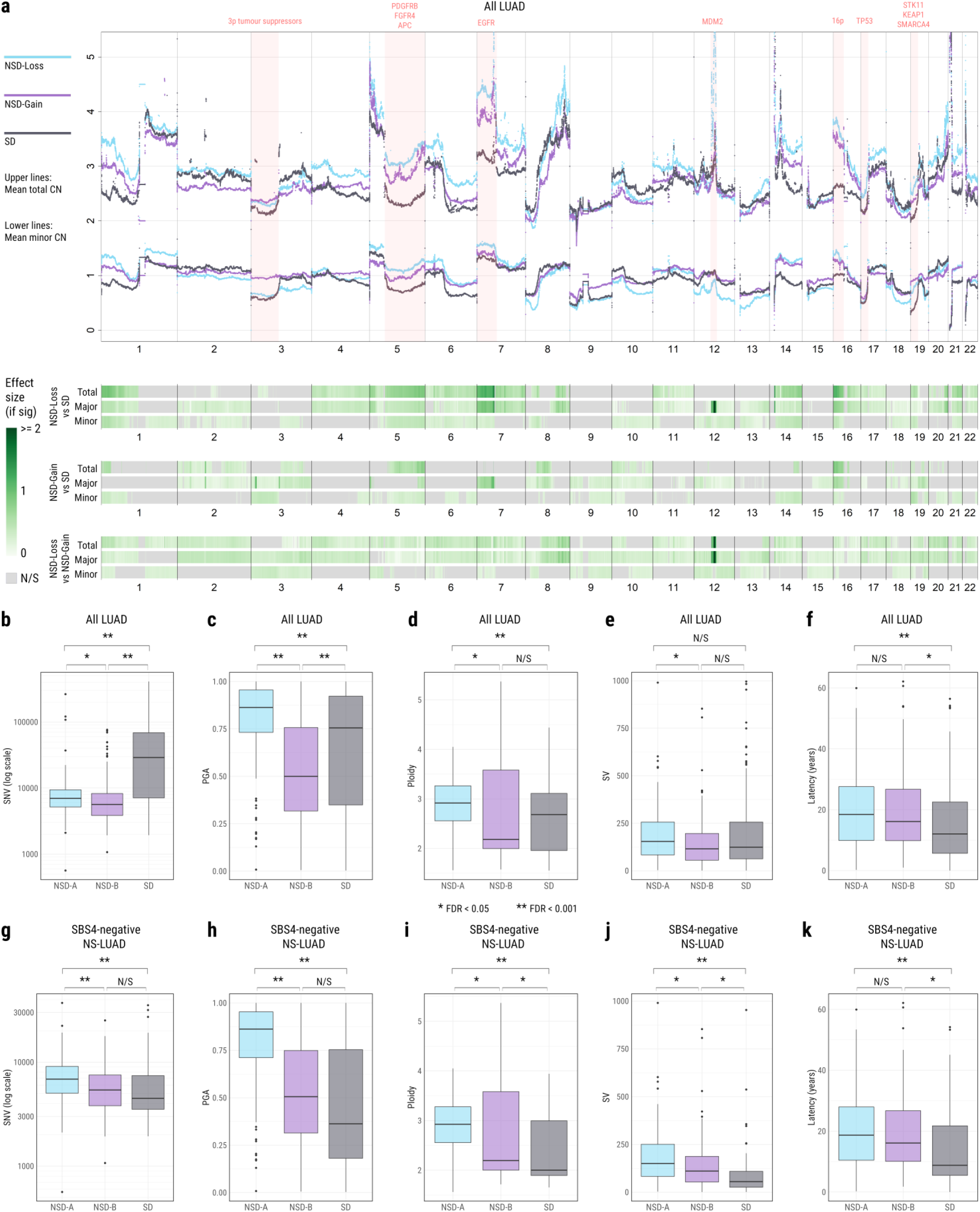
Comparison of overall states of tumours on different trajectories at the time of sampling. **a)** Mean nucleotide-level total and minor copy number of tumours on each trajectory. Blue lines represent *NSD-Loss*, purple lines represent *NSD-Gain*, and black lines represent *SD*. Upper lines in each colour represent total copy number. Lower lines in each colour represent minor copy number. Regions harbouring selected driver genes are highlighted in pink. Tracks below the copy number plot show loci with significant differences in total, major, and minor copy number between *NSD-Loss* & *SD* (top tracks), *NSD-Gain* & *SD* (middle tracks), and *NSD-Loss* & *NSD-Gain* (bottom tracks). Non-significant regions and those supported by fewer than 25% of samples in each class are marked grey. Regions with a significant difference are coloured by effect size, with a darker green representing a larger difference. **b-f)** Boxplots showing the comparisons between trajectories, across all LUAD, of **b)** SNVs, **c)** PGA, **d)** ploidy, **e)** SVs, and **f)** latency calculated with 1x acceleration. **g-k)** Boxplots showing the comparisons between trajectories, specifically within NS-LUAD, of **g)** SNVs, **h)** PGA, **i)** ploidy, **j)** SVs, and **k)** latency calculated with 1x acceleration. Midlines of boxplots represent median values. The coloured regions of each box represent the interquartile range (IQR) i.e. 25th - 75th percentile. Upper whiskers extend to the highest point not greater than the 75th percentile + 1.5xIQR. Lower whiskers extend to the lowest point not less than 25th percentile - 1.5xIQR. Outliers beyond these values are identified as dots.

One region stands out distinctly: a focal amplification on 12q centred on *MDM2*, showing markedly higher total and major copy number in NSD-Loss compared to NSD-Gain and SD tumours. *MDM2* is a well-established oncogene amplified in several tumour types^54–56^ including lung cancer^48,57^. Gains across the *MDM2* locus occur in similar proportions of samples across trajectories (**Tables S4-S7**). However, the amplification magnitude is significantly higher in NSD-Loss tumours than in NSD-Gain (**Fig. S5**, FDR = 0.0007) and SD tumours (FDR = 0.0001), indicating that amplification level, rather than occurrence frequency distinguishes this locus. While the major copy number is substantially increased at this locus, the minor copy number is not, pointing to this being an allele-specific amplification. Our recent work examined the role of extrachromosomal DNA (ecDNA) in lung cancer and found enrichment of MDM2 amplification via ecDNA in LCINS^58^. Here, NSD-Loss tumours are significantly more likely to undergo ecDNA amplification at the MDM2 locus than SD tumours (**Methods**, **Table S13**, risk ratio = 5.83, FDR = 0.029).

Across much of the genome, however, the copy number profiles of the three subsets follow broadly the same patterns; ploidy is gained and lost at the same loci. While there are statistically significant differences, the effect size at most loci is small: The difference in total copy number between NSD-Loss and NSD-Gain tumours is less than 0.5 (i.e. half a chromosome copy) for 94% of the genome. Between NSD-Gain and SD tumours, this is the case for 88% of the genome, and between NSD-Loss and SD tumours, it is the case for 74% of the genome. Two different routes—WGD tempered by losses, versus copy number gains from a diploid base—lead to similar overall copy number states. This implies that there may be broad selective pressures acting on LUAD tumours to reach a relative dose of oncogenes to tumour suppressors.

### NS-LUAD tumours on the SD trajectory show rapid evolution

Across all LUAD, SD tumours contain by far the greatest number of SNVs (**Fig. 4b**, **Table S14**, FDR vs NSD-Loss = 4.5×10^-16^, vs NSD-Gain = 3.6×10^-19^), as we would expect given the proportion of S-LUAD^9–11^. Meanwhile, NSD-Loss tumours display a significantly higher proportion of the genome altered (PGA) by SCNAs than the other trajectories (**Fig. 4c**, FDR vs NSD-Gain = 1.1×10^-19^, vs SD = 4.3×10^-6^). This is likely due to the mechanism of WGD combined with early losses and later gains - WGD samples have been shown to have higher PGA across tumour types^59^. Differences in ploidy are observed to be less pronounced than those in PGA (**Fig. 4d**), but again the NSD-Loss tumours display significantly higher ploidy than either NSD-Gain or SD tumours.

Surprisingly, NS-LUAD tumours on the SD trajectory display fewer genomic alterations than those on the NSD trajectories. When we consider the NS-LUAD tumours, those on the SD trajectory carry significantly fewer SNVs than NSD-Loss tumours (**Fig. 4g**, **Table S15**, FDR = 4.8×10^-4^), significantly lower PGA than NSD-Loss tumours (**Fig. 4h**, FDR = 7.1×10^-12^), and significantly lower ploidy than either NSD trajectory (**Fig. 4i**, FDR vs NSD-Loss = 2.9×10^-5^, vs NSD-Gain = 0.012). In fact, the median NS-LUAD tumour on the SD trajectory has ploidy of just 2.00.

When considering all LUAD, tumours on the NSD-Loss trajectory contain more structural variants than those on the NSD-Gain trajectory (**Fig. 4e**, FDR = 0.0025), but no significant differences are observed in the number of SVs in SD trajectory tumours compared to either NSD trajectory (**Fig. 4e**). In contrast, among NS-LUAD specifically, tumours on the SD trajectory contain the fewest SVs (**Fig. 4j**, FDR vs NSD-Loss = 4.8×10^-9^, vs NSD-Gain = 0.0021), and NSD-Loss tumours contain the most SVs (**Fig. 4j**, FDR vs NSD-Gain = 0.0029), in line with our companion article^15^. Yang et al. report *EGFR* mutations (here most strongly associated with NSD-Loss tumours and least common in SD tumours) to be positively associated with almost all types of complex SVs, whereas *KRAS* mutations (here most strongly associated with SD tumours and never observed in NSD-Loss tumours) are reported to be negatively associated with all types of simple SVs. When comparing the presence or absence of specific signatures between the trajectories, we find SV4 (clustered complex translocations) and SV6 (a mix of very large complex rearrangements) to be associated with NSD trajectories, both in all LUAD (**Fig. S6a**, **Table S9**, SV4 FDR = 4.7×10^-10^, SV6 FDR = 8.4×10^-6^) and among NS-LUAD (**Fig. S6b**, **Table S10**, SV4 FDR = 1.9×10^-4^, SV6 FDR = 0.0027). Meanwhile, SV5 (very short non-clustered deletions) is associated with the SD trajectory both in all LUAD (**Fig. S6a**, FDR = 1.2×10^-21^) and among NS-LUAD (**Fig. S6b**, FDR = 1.4×10^-4^).

Notably, SD tumours have shorter latency (time since the appearance of the most recent common ancestor of each tumour) than those on either NSD trajectory (**Methods, Fig. 4f**, **Table S14**, all LUAD FDR SD vs NSD-Loss = 1.1×10^-4^, vs NSD-Gain = 0.0043). The effect size of this difference is particularly accentuated in NS-LUAD tumours (**Fig. 4k**, **Table S15**, FDR SD vs NSD-Loss = 4.9×10^-4^, vs NSD-Gain = 0.0092). In fact, the median latency of NS-LUAD tumours on the SD trajectory is estimated to be just 8.78 years—approximately half that of NS-LUAD tumours on either NSD trajectory (NSD-Loss: 18.67 years, NSD-Gain: 16.09 years). Taken together with the enrichment for *EGFR* and *KRAS* mutations in the NSD and SD trajectories respectively, this finding is in agreement with our previous work showing shorter latency in KRAS mutant tumours and longer latency in *EGFR* mutant tumours, both of which were found to be stronger associations in NS-LUAD^8^. The fact that SD tumours progress to the point of diagnosis substantially faster than NSD tumours may partially explain the lower number of somatic alterations in NS-LUAD tumours on the SD trajectory: this more rapid evolution allows less time to accumulate alterations.

Despite these evolutionary differences, and S-LUAD subjects having poorer survival in comparison to NS-LUAD subjects (**Fig. S7c**), no significant differences are observed in survival between the trajectories. This is the case both when considering all LUAD (**Fig. S7a**) and specifically within NS-LUAD (**Fig. S7b**) after adjusting for age, sex, and tumour stage, and after applying multiple testing correction (**Methods**).

### Clonal *EGFR*, *KRAS* & *STK11* drive trajectory selection

Within each sample, each event was identified as occurring either clonally or subclonally. Events showing a significant preference between these states, compared to the average event within each trajectory, were then identified (**Methods**). Events that tend to occur clonally on a particular trajectory are more likely to be involved in driving a tumour toward that trajectory. On the other hand, events occurring preferentially subclonally are more likely to be a consequence of being on a given trajectory.

Mutations of *TP53* are preferentially clonal in all subsets (**Fig. 5**), highlighting its importance in initiating the disease, but not in fixing the trajectory a tumour will follow. *EGFR* mutations are preferentially clonal in both NSD trajectories (**Fig. 5a-b**). Conversely, *KRAS* and *STK11* mutations, as well as the LOH on 19p (*STK11*, *KEAP1*, *SMARCA4*) are preferentially clonal only in the SD trajectory (**Fig. 5c**). The occurrence of these events prior to subclonal expansion, specifically within these trajectories, suggests that they play an important role in committing a tumour to either the NSD trajectories or the SD trajectory. Notably, 57 of 58 (98%) of *STK11* mutant tumours also feature the LOH on 19p (**Fig. S2**, **Table S3**), implying that bi-allelic loss of *STK11* is a strong early driving mechanism in SD tumours.

**Fig. 5:**
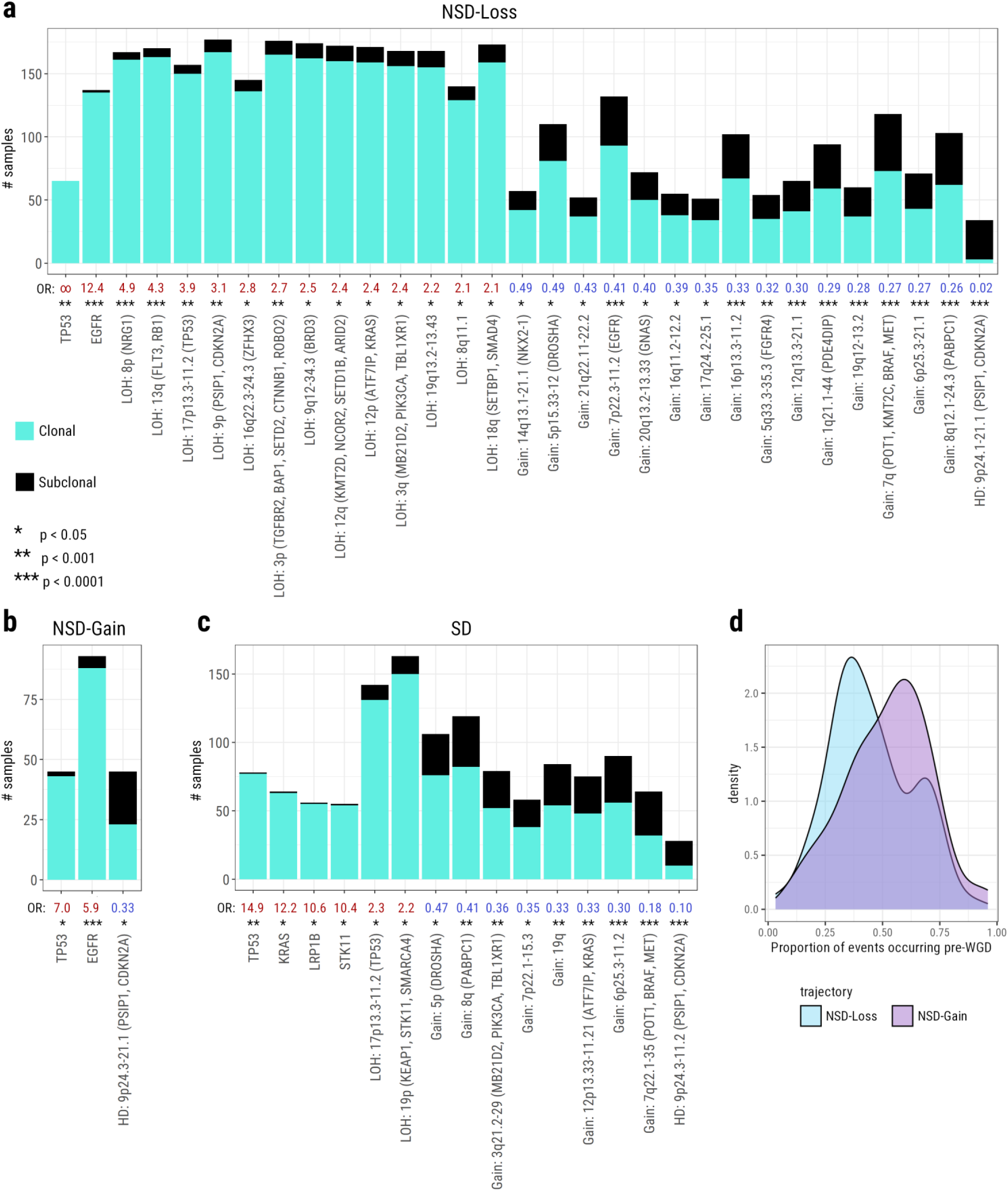
Clonal and subclonal preferences among key events within each trajectory. a-c) Barplots showing the number of clonal and subclonal occurrences of events with a significant preference, compared to all other events, to occur clonally or subclonally, within the **a)** NSD-Loss, **b)** NSD-Gain, and **c)** SD trajectories. Clonal occurrence count of each event is shown in cyan. Subclonal occurrence count is shown in black. Odds ratios of each event are shown below the bar plots. Odds ratios greater than 1 represent a preference for occurring clonally, and are shown in red. Odds ratios less than 1 represent a preference for occurring subclonally, and are shown in blue. Significance is shown below the odds ratios. A single asterisk represents p < 0.05. Two asterisks represent p < 0.001. Three asterisks represent p < 0.0001. Events without a significant preference for either clonality or subclonality are omitted from the figure. **d)** Overlaid density distributions of the proportion of events observed per sample that are ordered prior to WGD in NSD-Loss tumours (blue) and NSD-Gain tumours (purple). A Wilcoxon test indicates that the distributions are significantly different (p=0.009).

In the NSD-Loss trajectory, we observe that many LOH events exhibit a preference to occur clonally (**Fig. 5a**). This is not the case for any copy number events in NSD-Gain (**Fig. 5b**). This activity is likely linked to the more frequent, and earlier, WGD in NSD-Loss compared to NSD-Gain. Indeed, a significantly smaller proportion of key events occur prior to WGD in NSD-Loss tumours than in NSD-Gain tumours (**Fig. 5d**, *p* = 0.009). This further elucidates the evolutionary distinction between the two NSD paths; when *EGFR* mutations are coupled with early WGD and substantial clonal copy number losses, this generally pushes tumours to the NSD-Loss trajectory.

In both NSD-Loss and SD we observe several copy number gains occurring subclonally more often than the average event (**Figs. 5a & 5c**). This pattern is not observed in NSD-Gain (**Fig. 5b**). This reiterates our earlier finding that the patterns of copy number alterations are similar in NSD-Loss and SD tumours, with losses generally preceding gains, while losses are interspersed among the gains in NSD-Gain tumours.

Mutations of *EGFR*, *KRAS*, and *STK11* are the events that show the strongest associations with either the NSD trajectories or the SD trajectory (**Fig. 3**). They are also among the most consistently early events on their trajectories (**Figs. 2 & 5**). This implies that a tumour’s trajectory is often fixed early in its progression. However, if we consider NS-LUAD samples without any of these mutations, we find that the loss of 3p remains significantly associated with the SD trajectory, whereas gains of 7p (*EGFR*) and both arms of chromosome 16 remain significantly associated with the NSD trajectories (**Fig. S8**, **Table S17**). These events occur further down the respective evolutionary trajectories (**Fig. 2**), implying that - in the absence of *EGFR*, *KRAS*, or *STK11* mutations - these events may influence later fixation to an evolutionary trajectory.

### *KRAS* allows NS-LUAD tumours to progress without genomic instability

In order to assess each mutation’s impact on the evolution of copy number burden, we calculated the proportion of copy number signature activity^60,61^ classed as aberrant (**Methods**). Despite some associations, e.g. between smoking status, *EGFR*, and *KRAS*, our data did not show high levels of multicollinearity^62,63^(**Fig. S9**), allowing us to perform multivariate linear analysis (**Methods**).

Our analysis reveals that *EGFR* and *TP53* mutations are associated with more aberrant copy number activity, i.e. greater genomic instability (**Fig. 6a-b**; p = 4.4×10^-7^ and p = 4.8×10^-8^ respectively). *EGFR* mutant tumours deviate from their baseline copy number state significantly more than *EGFR* wild-type tumours (**Methods**, **Figs. 6b-c**; FDR = 7.4×10^-5^). Conversely, *KRAS* mutations are associated with less aberrant copy number activity, i.e. more stable genomes (**Figs. 6a & 6d**; p = 7.9×10^-5^). *KRAS* mutant tumours deviate from their baseline copy number state significantly less than *KRAS* wild-type tumours (**Figs. 6d-e**, FDR = 0.0014). Additionally, *KRAS* mutation is negatively associated with a number of copy number events (**Fig. S2**, **Table S3**). These findings imply that *EGFR* or *TP53* mutations could be driving genomic aberrations in the tumours in which they occur. However, KRAS-mutant tumours do not appear to require as much copy number activity to progress.

**Fig. 6:**
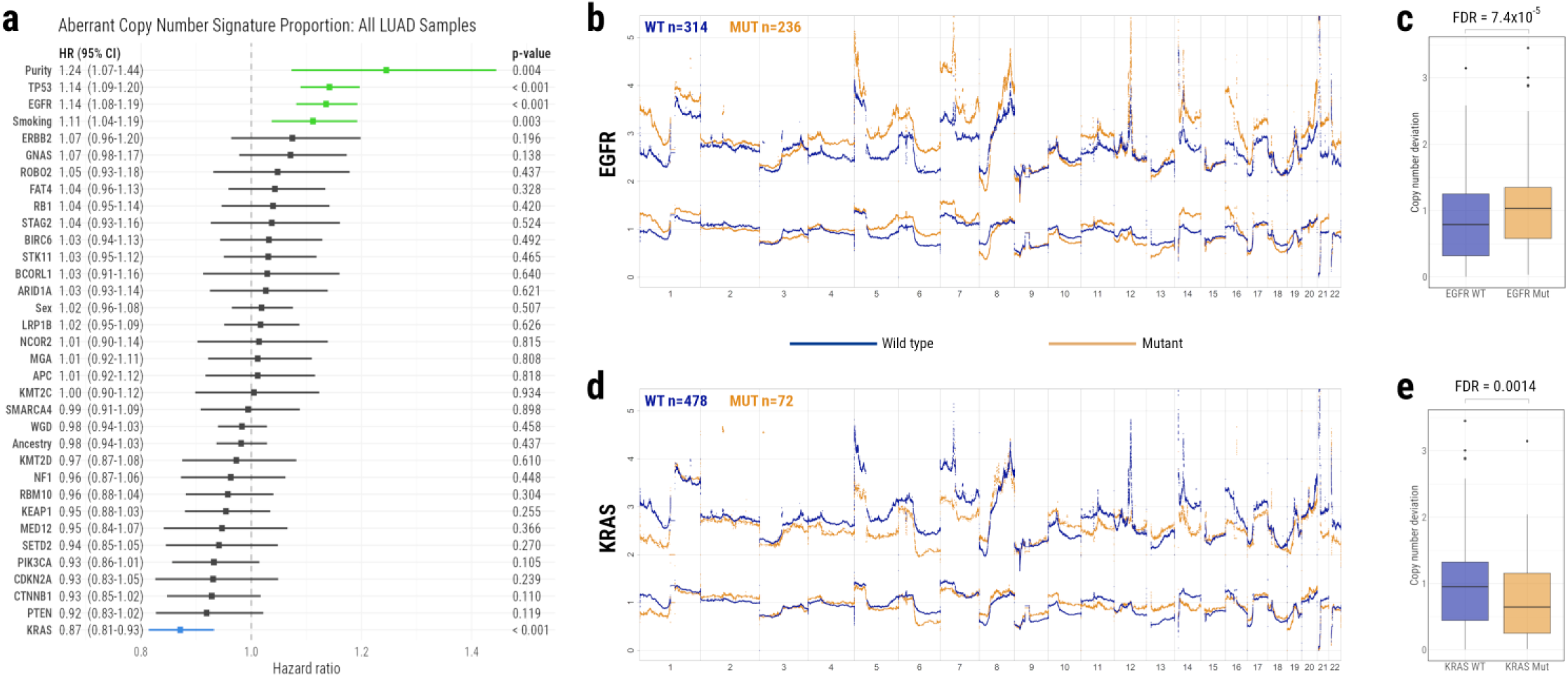
The association between driver gene mutations and genomic instability. **a)** Forest plot of multivariate linear analysis investigating the association between mutations in all identified driver genes, and the proportion of copy number signature activity that was aberrant. Aberrant copy number signature activity refers to signatures other than CN1 (diploidy) in non-WGD samples, and signatures other than CN2 (tetraploidy) in WGD samples. Driver genes, WGD, and potentially associated demographic features (tumour purity, smoking status, sex, ancestry) are ordered by hazard ratio. Hazard ratios with confidence intervals greater than 1 represent significant associations with more aberrant copy number signature activity. These are coloured in green. Hazard ratios with confidence intervals less than 1 represent significant associations with less aberrant copy number signature activity. These are coloured in blue. **b-c)** Average copy number profiles of tumours that have wild type vs mutant **b)** EGFR and **c)** KRAS. The copy number of wild type samples is shown in blue. The copy number of mutant samples is shown in orange. Upper lines represent total copy number. Lower lines represent minor copy number.

Further, this finding provides an explanation as to why NS-LUAD tumours on the SD trajectory have fewer genomic alterations than those on the NSD trajectories: these tumours are significantly enriched for *KRAS* mutations. Without the added impact of tobacco smoking, these tumours have quieter genomes than their *EGFR* counterparts. In NS-LUAD tumours, *EGFR* mutations occur exclusively on the NSD trajectories.

## Discussion

Our *de novo* event-ordering analysis discovered three distinct evolutionary trajectories in LUAD. Two of these—NSD-Loss and NSD-Gain—are dominant in NS-LUAD tumours, while the third trajectory—SD—is followed by most S-LUAD tumours. Notably, 18% of NS-LUAD tumours are observed to follow the smoking-like SD trajectory. In addition to smoking status, genetic ancestry was found to be associated with NS-LUAD tumour evolution, with individuals of European ancestry approximately three times more likely than those of East Asian ancestry to develop tumours with the SD trajectory. In contrast, passive smoking was not found to have any impact on the likelihood of a tumour taking the SD trajectory, aligning with our previous findings of no association between passive smoking and mutational signatures or driver gene alterations^6^. Each evolutionary trajectory is defined by distinct genomic events. Both NSD trajectories feature early *EGFR* mutations. NSD-Loss is further characterised by a frequent WGD tempered by copy number losses, while NSD-Gain usually gains ploidy via copy number gains from a diploid base. The SD trajectory is largely driven by driver mutations in *KRAS* and *STK11*.

We observed a degree of convergence in genome-wide copy number states, suggesting overarching selective pressures to reach a relative dose of oncogenes to tumour suppressors. However, the targeting of distinct genes on different trajectories implies varying pathway activity. The MAPK pathway is implicated in the SD trajectory due to the importance of *KRAS* mutations^16–18,20–25^. Meanwhile in NSD tumours, *EGFR* mutations and copy number gains, in addition to the gains of additional growth factor receptor genes *PDGFRB* and *FGFR4*, suggest that this pathway is activated via a different mechanism^17,20,21,25,32^. Alternatively, other *EGFR*-related pathways may be activated^25,32,33^. Additionally, 3p tumour suppressors have been previously reported as being lost more frequently in S-LUAD than NS-LUAD^30,31^. Our analysis reveals that this loss is still a common event in NSD-Loss tumours, and is rare specifically in NSD-Gain tumours.

Interestingly, NS-LUAD tumours of the SD trajectory display quieter genomic activity compared to those on the NSD trajectories; They lack the increased mutational burden that is induced by tobacco smoking^9^, and have significantly lower PGA and ploidy than NS-LUAD tumours on the NSD trajectories. This is explained by the strong enrichment for *KRAS* mutations among NS-LUAD tumours on the SD trajectory, versus the enrichment for *EGFR* mutations among those on the NSD trajectories. *EGFR* mutant tumours were shown to have increased genomic instability. Conversely, *KRAS* mutant tumours were shown to have more stable genomes, implying that they do not require as many copy number changes to progress. These findings are supported by our companion article which reports higher SV burden in *EGFR*-mutant tumours and lower SV burden in *KRAS*-mutant tumours^15^. The reduced genomic instability in NS-LUAD tumours on the SD trajectory may be at least partially due to their short latency; more rapid evolution allows less time to accumulate SCNAs.

Among people who have smoked, LUAD tumours in patients of European genetic ancestry have previously been reported to exhibit greater genomic instability than those of East Asian genetic ancestry, while no significant difference was reported among people who have never smoked^64^. Further, specific SCNAs affecting 16p and 19p have been reported as enriched in LUAD tumours in people from East Asia and Western Europe respectively^65^. Meanwhile, LUAD tumours in people who have never smoked have been shown to display more extensive SCNAs than in those who have smoked^66^. Using evolutionary orderings, we have incorporated these previous observations into a unified model, in which 16p and 19p SCNAs occur predominantly in NS-LUAD and S-LUAD trajectories, respectively, and tumours in the SD trajectory have lower levels of genomic instability. Further, the data-driven approach taken in this study has allowed us to go beyond manual subsetting on the basis of demographic information and look more deeply at the underlying genomic events that define tumour evolution. This allows a more personalised understanding of how an individual’s tumour is evolving, and how it may be expected to continue to evolve.

Our previous research, based on a subset (n=232 NS-LUAD tumours) of our current cohort, classified tumours into one of three manually-curated groups based on copy number profiles, i.e. snapshots of the overall state of each tumour at the time of sampling^4^. That research was able to hint at some of the findings in our current study. For example, the quietest manually-curated NS-LUAD subset—“*Piano*”—featured the highest rate of *KRAS* mutations and the lowest rate of *EGFR* mutations. This study leverages a larger cohort of both NS-LUAD and S-LUAD tumours, and utilises an event-based approach: tumours are classified based on a broader range of genomic information including both SNVs and SCNAs, and factoring in the order in which these events are observed. This leads to a starker separation between *EGFR*-mutant and *KRAS*-mutant tumours, while also highlighting the association of other events such as *STK11* mutation in the determination of which evolutionary trajectory a tumour will follow. This has enabled us to uncover and explore in depth the underlying dynamics of the varying modes of evolution associated with these alterations. Importantly, our *de novo* discovery of these trajectories enabled us to identify a subset of NS-LUAD tumours evolving in a smoking-like manner.

Together, our findings demonstrate that although NS-LUAD tumours generally lack the high mutational burden observed in S-LUAD, a substantial subset - one in six - follow a smoking-like genomic trajectory. As these are commonly *KRAS* mutant tumours, they require fewer copy number changes than their *EGFR* mutant counterparts to progress. The emergence of *KRAS* inhibitors in recent years has provided hope for improved treatment of *KRAS*-mutant lung cancer^67–73^. Another common treatment option is the use of immune checkpoint inhibitors (ICIs)^74,75^. However, immunotherapy is not always effective^76–78^, and can result in increased toxicity and immune-related adverse events (irAEs)^79,80^. High TMB is considered a response biomarker for immunotherapy in lung cancer, among other tumour types^81–84^. Many NS-LUAD tumours on the SD trajectory harbour *KRAS* mutations, but do not have an elevated mutation burden overall. As such, while these patients may be responsive to *KRAS* inhibitors, they may be less likely to respond to immunotherapy. Careful consideration of these factors should be given to each individual when aiming to optimise treatment efficacy while minimising unnecessary side effects. The distinct evolutionary trajectories presented in this study could therefore have critical implications for prognosis, molecular classification, and therapeutic targeting in lung cancer from people who have never smoked.

## Methods

### Dataset & Quality Control

The Sherlock-Lung study contained a total of 1217 whole genome sequencing (WGS) tumour samples, with matched normal pairs, at the time of this study. 1032 of these were LUAD tumours. It was deemed to be important to analyse a single histology in order to avoid *de novo* subsetting being biased by histological differences. LUAD were selected as they represented 85% of the overall cohort and had a specific genomic landscape^6,8^. Quality control was carried out on the outputs of Battenberg copy number calling and DPClust mutational subclonal clustering. Samples were refitted with further runs of Battenberg and DPClust if they were deemed to have been assigned incorrect purity or ploidy. As many rounds of refitting as necessary could be employed to find a plausible solution for each sample. Samples were included in this study if three criteria were met: (1) A minimum number of reads per chromosome copy (NRPCC) of 10, (2) A minimum purity of 30%, (3) Biologically plausible outputs from Battenberg and DPClust. We determined that these criteria provided the best chance for us to identify subclonal expansions, where they occur. In fact, all samples that passed criteria (1) and (2) did have plausible Battenberg and DPClust outputs, after necessary refits. This resulted in a set of 550 LUAD tumours, of which 407 had never smoked and 143 had smoked.

### Ancestry estimation

The genetic ancestry of each patient was assigned based on genetic similarity to labelled reference superpopulations from the 1000 Genomes Project^85^, using VerifyBamID version 2.0.1^86^ and normal-tissue WGS data.

### Ordering events in de novo subsets of samples

The *de novo* discovery of subsets of samples, and the ordering of events within those subsets, was carried out via a Plackett-Luce ordering model.

#### i. Identifying commonly occurring somatic copy number alterations

Copy number profiles were produced for each sample using Battenberg. Based on these profiles, segments were called as LOH, non-LOH loss, HD, gain, or no SCNA by considering their copy number and whether the sample had undergone WGD. For example, a segment with copy number 2+1 would be classed as a gain in a non-WGD sample, but a non-LOH loss in a WGD sample. Binomial probability was used to identify loci at which gains, LOH, or HDs occurred in more samples than expected by chance. To obtain only the most certain and robust events, regions were filtered out if they occurred at any of the following loci:

- Centromeres
- Within 5 megabases of the telomeres
- The human leukocyte antigen (HLA) region
- Short arms of acrocentric chromosomes 13, 14, 15, 21, and 22 - In particular, we confirmed the identification of copy number gains on 21p and 22p in a high proportion of samples as reported in our previous work^8^. However, for the purpose of this study these were conservatively filtered out due to the possibility of identifying events artefactually.
- X and Y chromosomes - as the NS-LUAD and S-LUAD cohorts in this study contained vastly different numbers of male and female subjects, we did not wish to bias de novo discovery by using events on these chromosomes.

Overlapping regions were merged across all samples to generate a population-wide set of commonly occurring SCNAs.

#### ii. Ordering events within individual samples

The identified SCNA segments, as well as SNVs in a pre-specified list of potential driver genes^6,87^ occurring in at least 3% of samples in the set, were ordered within each sample. Each event was identified as either clonal or subclonal (based on Battenberg outputs for SCNA segments and DPClust outputs for SNVs). Clonal events were ordered before subclonal ones. In WGD samples, clonal events were sub-classified as “early clonal” (pre-WGD) or “late clonal” (post-WGD) based on their copy number status. For example, in a WGD sample, a segment with clonal copy number 2+0 would be called as an early clonal LOH, whereas a segment with clonal copy number 2+1 would be called as a late clonal non-LOH loss. Early clonal events were ordered before late clonal ones. Subclonal events were ordered according to a phylogenetic tree, which was generated according to the events’ CCFs and was required to satisfy the pigeonhole principle. In the case of multiple phylogenetic trees being equally likely, one was selected at random. In cases where events could not be definitively ordered relative to each other within a sample (e.g. different early clonal events, or events in branching subclones), these events were ordered randomly relative to one another. In order to account for the stochasticity in this aspect of the ordering, and the selection of phylogenetic trees, the ordering was repeated for 1000 iterations.

#### iii. *De novo* discovery of subsets of samples

The PLMIX R package^88^ was used to generate mixture models, identifying a specified number of subsets of different event trajectories. This was run to generate sets of 1, 2, 3, 4, and 5 subgroups. The number of subgroups resulting in the lowest median Bayesian information criterion score was selected as the optimal number of subgroups.

#### iv. Repeat steps i and ii for each subset

Steps i-iii were run across a whole set of samples, to perform *de novo* identification of subgroups. For each subgroup, steps i and ii were then re-run to ensure that any subset-specific events could be identified and included in the subgroup’s trajectory.

#### v. Calculate and plot aggregate ordering for each subset

With an order established within each individual sample, an overall order was collated across all samples in each subgroup using the PlackettLuce R package^89^. Median values with 95% confidence intervals are plotted.

### Demographic comparisons

To reduce the effect of confounding variables, comparisons of demographic splits between the trajectories were performed on subsets of the data. The comparison of self-reported smoking statuses was performed on all LUAD, and additionally performed specifically on tumours in European individuals in order to control for the potential confounding effect of ancestry; European was the largest ancestry group, and the second-largest ancestry group (East Asian) contained only two subjects who reported having smoked. The comparison of ancestry was performed on tumours in people who report they have never smoked - as the largest smoking status group - and between European and East Asian ancestry - as the two largest ancestry groups. The SBS4 and passive smoking comparisons were performed on tumours in people who report they have never smoked. The sex comparison was performed on tumours from people with both smoking statuses, as males accounted for most of those who have smoked and females accounted for most those who have never smoked in our data. For each comparison, the proportion of tumours from each subgroup (e.g. males, females) assigned to each trajectory were plotted as a stacked percentage bar graph. The number of tumours in each subgroup assigned to the SD trajectory, and the two NSD trajectories combined, were collated in a confusion matrix. Odds ratios and risk ratios were calculated based on these confusion matrices. A two-tailed Fisher exact test was performed on the confusion matrix for each comparison. Multiple testing correction was carried out on the resultant p-values using the Benjamini-Hochberg method to calculate false discovery rates.

### Comparing the number of driver genes with SNVs identified in >= 3% of samples across trajectories

A confusion matrix was calculated containing the number of driver genes from a predefined list^6,87^ (n=72) that did, vs did not, contain SNVs in >= 3% of samples in each subset of tumours (NSD-Loss, NSD-Gain, SD). For each trajectory, a Fisher’s exact test was performed to check for a significant difference between the number of driver genes identified in the given subset vs the other two subsets combined. Multiple testing correction was carried out on the resultant p-values using the Benjamini-Hochberg method to calculate false discovery rates.

### Event co-occurrence and mutual exclusivity

Positive and negative associations between pairs of events, across all LUAD, were calculated using hypergeometric distribution, via the *cooccur* R package^90,91^. Multiple testing correction was carried out on the p-values of each set of associations (positive, negative) using the Benjamini-Hochberg method to calculate false discovery rates. Events with at least one significant FDR for association with another event were included in the figure.

### Event-level comparisons

In order to compare like-for-like events across different trajectories, the set of events used was that which was identified across all samples during the initial *de novo* discovery phase of the ordering algorithm. For each event, a confusion matrix was generated stating the number of samples in which each event was present and absent in each subset (e.g. NSD-Loss vs NSD-Gain, or the SD trajectory vs combined NSD trajectories). A two-tailed Fisher test was applied to each event’s confusion matrix. Multiple testing correction was carried out on the resultant p-values using the Benjamini-Hochberg method to calculate false discovery rates. Volcano plots were generated, highlighting any events with a significant (FDR < 0.05) and substantial (odds ratio > 3/2 or < 2/3) association with one trajectory, or set of trajectories, within a given comparison.

### Mutational signature analysis

*De novo* mutational signatures for single base substitutions (SBSs), doublet base substitutions (DBSs), indels (IDs), copy number alterations (CNs), and structural variants (SVs) were extracted using SigProfilerExtractor^92^ v1.1.21 with default parameters, normalization set to 10,000 mutations (only for SBSs, DBSs, and IDs), and splitting the cohort of 1,217 lung cancer patients between people who have smoked (*n* = 345) and people who have never smoked or have unknown smoking status (*n* = 872). SBS *de novo* signatures were extracted using the SBS-288 and SBS-1536 high-definition mutational contexts, which, beyond the common SBS-96 trinucleotide context using the mutated base and the 5’ and 3’ adjacent nucleotides^93,94^, also consider the transcriptional strand bias and the pentanucleotide context (two 5’ and 3’ adjacent nucleotides), respectively^94^. Given the high similarity obtained for both mutational contexts, as well as the additional separation of mutational processes obtained for the SBS-288 mutational context, the SBS-288 context results were used for further analysis. The previously established mutational contexts DBS-78 and ID-83 were used for the extraction of DBS and ID signatures^94,95^. CN signatures were extracted *de novo* following an updated context definition benefitting from deep WGS data (CN-68), which allowed further characterization of CN segments of less than 100 kbp in length (in contrast to current COSMICv3.4 reference signatures using the CN-48 context, which are based on SNP6 microarray data, and therefore lack the resolution to characterize short CN segments)^60^. SV signatures were extracted using a similarly refined context, with an in-depth characterization of short SV alterations below 1 kbp (SV-38). After *de novo* extraction was completed, SigProfilerAssignment^96^ v0.1.1 was used to decompose the *de novo* extracted mutational signatures into COSMICv3.4^97^ reference signatures based on the GRCh38 reference genome, as well as to assign signatures to individual samples obtaining signature activities, based on the forward stagewise algorithm for sparse regression and nonnegative least squares for numerical optimization, as previously described^6,8^.

SBS, DBS, ID, CN, and SV signatures were assigned for all tumour samples as described above. Comparisons were then carried out calculating the preference of all signatures between the SD trajectory and both NSD trajectories combined, as described in the context of individual somatic events in the above section “*Event-level comparisons”*.

### Average copy number comparisons

Copy number segments were identified for each sample by Battenberg^98^. For a given set of samples, a combined set of segment loci was generated with segments delineated at every breakpoint in any sample in the set. The copy number state for each sample, in each of these segments, was then calculated. If the segment was subclonal, copy number states were reported as a weighted average of the different subclones, with weights being determined by the CCF (i.e. the proportion of cells in which a given copy number state is reported). The mean of the major, minor, and total copy number states were then calculated at each segment in each sample set. The distributions of the copy number states were compared, at each segment, for each pair of subsets (i.e. NSD-Loss vs SD, NSD-Gain vs SD, and NSD-Loss vs NSD-Gain), using Wilcoxon rank sum tests. Multiple testing correction was carried out on the resultant p-values using the Benjamini-Hochberg method.

### Subset-wide metric comparisons

For each tumour sample, the following metrics were calculated: number of SNVs, PGA, ploidy, number of SVs, kataegis level, latency, survival (in weeks), age at diagnosis, NRPCC, and tumour purity. Latency refers to the difference in years between estimated age of the patient at the emergence of the most recent common ancestor (MRCA) in tumour evolution and the age at diagnosis, assuming a constant mutation rate per year. This method has been utilised in our previous research^4,8^, and was described in the PCAWG evolutionary study^99^. The distributions of these metrics were compared between trajectories, pairwise (i.e. NSD-Loss vs SD, NSD-Loss vs NSD-Gain, and NSD-Gain vs SD), using Wilcoxon rank sum tests. Within each pairwise comparison, multiple testing was carried out across the resultant p-values for all metrics using the Benjamini-Hochberg method. These comparisons were performed considering all LUAD samples, as well as specifically among NS-LUAD, and among S-LUAD. Comparisons of SNVs, PGA, ploidy, SVs, and latency were visually compared using boxplots for all LUAD, and for NS-LUAD.

### Major copy number comparison at MDM2 locus

The number of samples in which copy number gains were present and absent at the MDM2 gene locus were calculated for each trajectory. Confusion matrices were generated for each pairwise comparison of trajectories (NSD-Loss vs NSD-Gain, NSD-Loss vs SD, and NSD-Gain vs SD). Two-tailed Fisher exact tests were performed on these confusion matrices, but showed no significant differences between the number of gains at this locus after multiple testing correction. The weighted average major copy number was calculated for each identified gain at the MDM2 locus by multiplying the major copy number of the gained segment by the size of the gain (capped at the size of the gene), and dividing by the size of the gene. These major copy numbers were plotted as boxplots, comparing the three trajectories. The major copy numbers were further compared by Wilcoxon rank sum tests between each pairwise comparison of trajectories. Multiple testing correction was carried out on the resultant p-values using the Benjamini-Hochberg method.

### ecDNA amplification comparison at the MDM2 locus

In order to identify ecDNA amplifications, CNVKit v.0.9.6^100^ was run in tumour-normal mode to call SCNAs, AmpliconArchitect v.1.3^101^ was used to reconstruct amplicon architecture, and AmpliconClassifier v.1.3.1 was used to classify the most probable mechanism by which each amplicon was formed, as described in detail in Khandekar et al.^58^. A confusion matrix was calculated indicating the number of tumour samples with and without an ecDNA amplification at the MDM2 locus for each subset of tumours (i.e. those following the NSD-Loss, NSD-Gain, and SD trajectories). Two-tailed Fisher exact tests were performed on the confusion matrix for each pairwise comparison between trajectories. Multiple testing correction was carried out on the resultant p-values using the Benjamini-Hochberg method to calculate false discovery rates.

### Event clonality categorisation and preferential timing analysis

Somatic events were divided into those which were clonal and those which were subclonal, according to their CCF. In order to analyse clonality preferences, a confusion matrix was generated for each event within each evolutionary trajectory. These matrices recorded the number of times the event occurred both clonally and subclonally, as well as the number of times all other events occurred both clonally and subclonally. Fisher tests were performed in order to identify events that statistically occurred preferentially clonally or preferentially subclonally. Multiple testing correction was carried out on the resultant p-values using the Benjamini-Hochberg method. This analysis was performed on all events combined (mutations, gains, losses, and homozygous deletions) in order to identify the events that occurred preferentially clonally/subclonally overall across the given evolutionary trajectory.

### Comparison of the proportion of events observed prior to WGD

For each sample in which WGD was observed, the number of events occurring prior to WGD was calculated. This was then converted into a proportion of the total number of identified events occurring in the sample. The distribution of these proportions was plotted for both NSD-Loss and NSD-Gain trajectory tumours to compare the ordering of WGD between the two. To compare statistically, a Wilcoxon rank sum test was performed, comparing the proportions of events occurring prior to WGD in NSD-Loss versus NSD-Gain samples.

### Survival analysis

Adjusted survival curves were plotted comparing the three trajectories across all LUAD, and specifically among NS-LUAD. Further adjusted survival curves were plotted comparing people who self-reported as having smoked with those who self-reported as never having smoked across all LUAD. For each curve, a Cox proportional hazards model was calculated including survival time in weeks, as well as confounders tumour stage, sex, and age. In the comparisons between trajectories, self-reported smoking status was included as an additional confounder. Adjusted curves were then calculated using the direct standardisation method^102^, based on these Cox proportional hazards models. In order to investigate statistical differences between survival of different groups, adjusted curve tests - modified Pepe and Flemming tests for the difference between two adjusted survival curves^102^ - were carried out at 5 years.

### Genomic instability analysis

#### i. Defining aberrant copy number activity

CN signatures were assigned as described in the above section “*Mutational signature analysis”*. Signature CN1 was considered the baseline signature in non-WGD samples, and CN2 was considered the baseline in WGD samples. All other signature activity was considered “aberrant”. The proportion of signature activity that was aberrant was calculated for each sample.

#### ii. Multivariate linear regression

A single multivariate linear regression model was fitted with the dependent variable being the above defined proportion of aberrant copy number signature activity, and the explanatory variables being the presence or absence of each putative driver gene and WGD, the tumour purity, self-reported smoking status, ancestry, and sex. The resultant model was displayed as a forest plot, with significant negative associations between aberrant copy number and a variable (i.e. greater genomic stability) highlighted in blue, and significant positive associations between aberrant copy number and a variable (i.e. greater genomic instability) highlighted in green.

#### iii. Multicollinearity testing

A numeric matrix was calculated, comprising the all variables in the multivariate linear regression model, with the exception of aberrant copy number level: tumour purity, and binary values representing self-reported smoking status (have smoked = 1, never smoked = 0), ancestry (European = 1, other = 0), sex (male = 1, female = 0), and the presence or absence of each putative driver gene and WGD (present = 1, absent = 0) for each sample. The correlation between each pair of variables in the numeric matrix was then calculated. The largest absolute correlation was that between sex and smoking, at 0.529, below the recommended threshold of 0.8 for variables to be considered collinear^62^. A correlation plot was produced, with positive correlations displayed in blue, and negative correlations displayed in red. Larger and darker circles represented higher absolute correlation. Variance inflation factor (VIF) values were also calculated for each variable in the multivariate linear regression model. These were plotted as a bar graph. The largest VIF value was that of smoking, at 2.41, below the recommended threshold of 5 for multicollinearity to be considered problematic for the interpretation of the linear model^62,63^.

#### iv. Copy number deviation

Copy number deviation from the baseline state was calculated as the mean distance from 1+1 in non-WGD samples, and distance from 2+2 in WGD samples. For example, a diploid tumour (baseline state 1+1) with average major copy number of 1.5 due to gains, and average minor copy number of 0.5 due to LOH, would have a deviation value of 1 (0.5 absolute major copy number deviation + 0.5 absolute minor copy number deviation). Differences between deviation scores among mutant and wild-type tumours were tested with Wilcoxon rank sum tests. Multiple testing correction was carried out on the resultant p-values using the Benjamini-Hochberg method to calculate false discovery rates.

## Supplementary Data

**Tables S1-S18**

See accompanying document

### Supplementary figure captions

**Figure S1:**
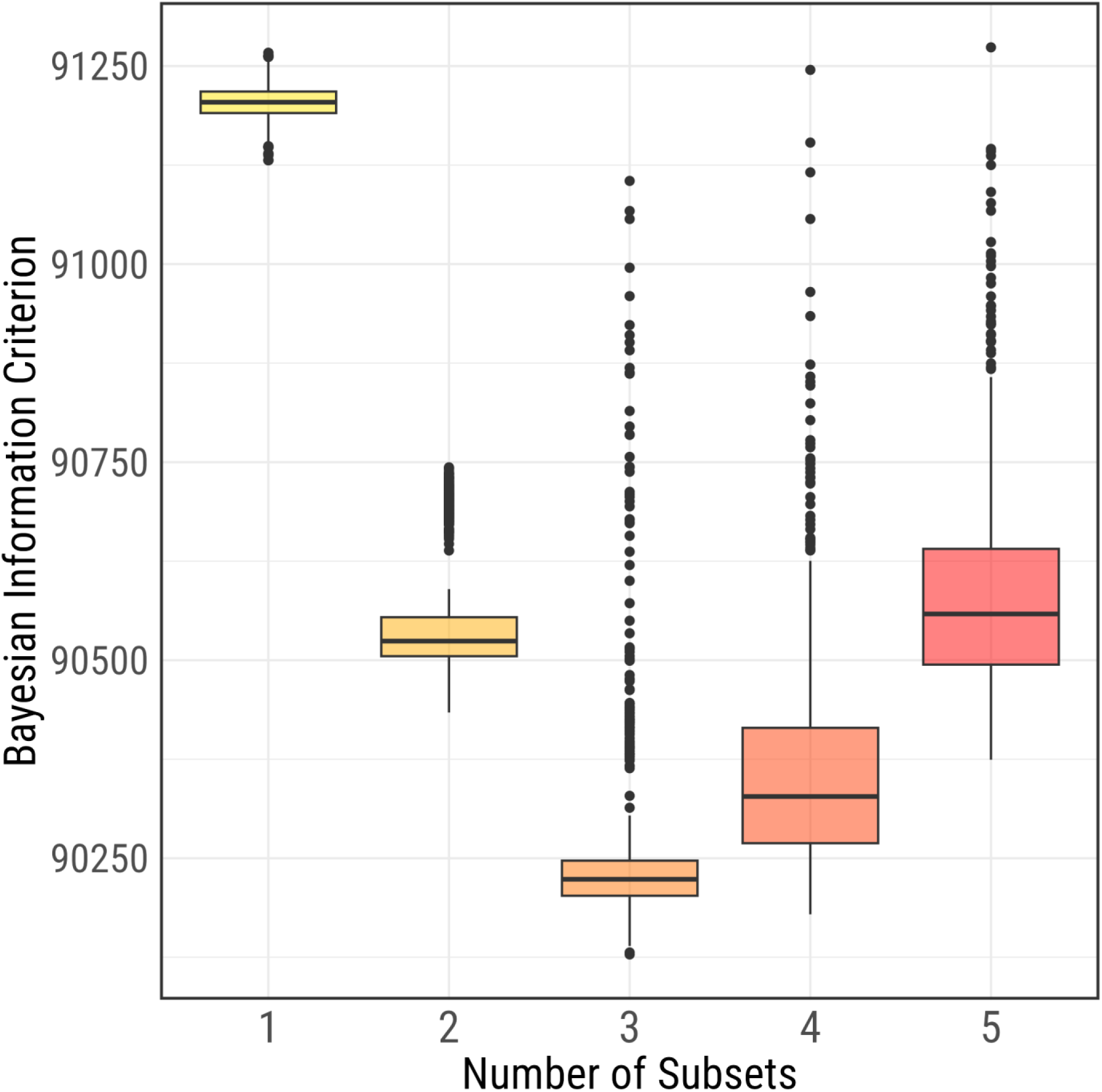
Selection of the optimal number of trajectories. Boxplots showing the Bayesian Information Criterion (BIC) scores for 1000 iterations with each number of subsets from 1 to 5. 3 subsets were selected as optimal due to having the lowest median (BIC) value. Wilcoxon rank sum tests were performed on each pairwise comparison between numbers of subsets. Multiple testing corrected FDRs were < 1×10^-6^ for each pairwise comparison.

**Figure S2:**
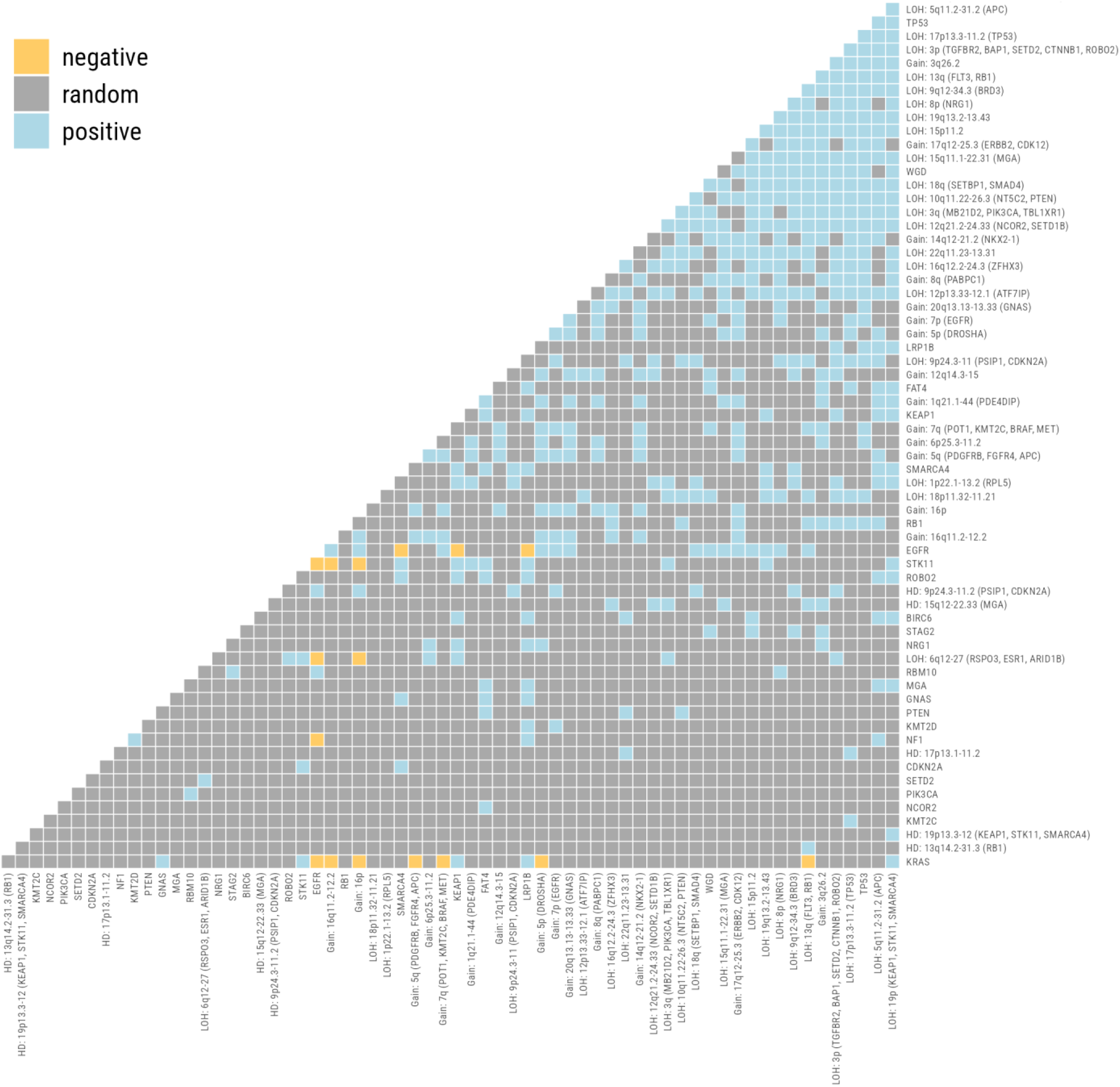
Event co-occurrence and mutual exclusivity. Positive and negative associations between pairs of events across all LUAD, as calculated by hypergeometric distribution. Positive associations are shown in blue, negative associations in yellow, and lack of significant association shown in grey. Events with no significant positive or negative associations with any other event are excluded from the figure.

**Figure S3:**
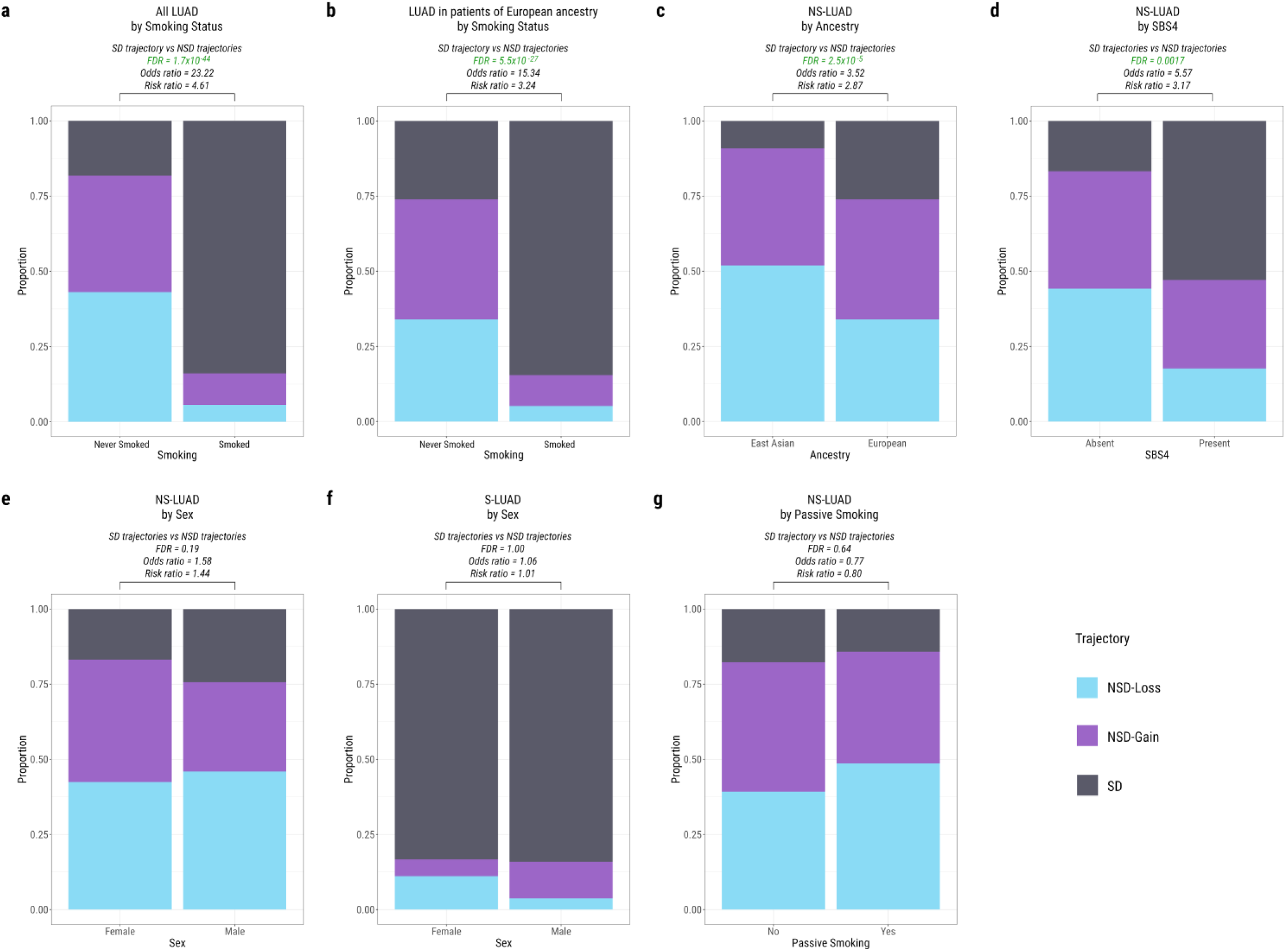
Demographic distribution across trajectories. Stacked bar plots showing the proportion of each demographic assigned to each trajectory. The proportion on the NSD-Loss trajectory is shown in blue, NSD-Gain in purple, and SD in grey. Statistical comparisons shown above barplots are two-tailed Fisher tests performed on the SD trajectory versus the combined NSD trajectories. Significant FDRs are written in green text. Comparisons shown are a) all LUAD split by self-reported smoking status, b) tumours in individuals of European ancestry split by self-reported smoking status (smoked vs never smoked), c) tumours in people who self-reported as having never smoked split by ancestry (European vs East Asian), d) tumours in people who self-reported as having never smoked split by SBS4 status (present vs absent), e) tumours in people who self-reported as having never smoked split by sex (male vs female), f) tumours in people who self-reported as having smoked split by sex (male vs female), and g) tumours in people who self-reported as having never smoked split by passive smoking exposure (yes vs no).

**Figure S4:**
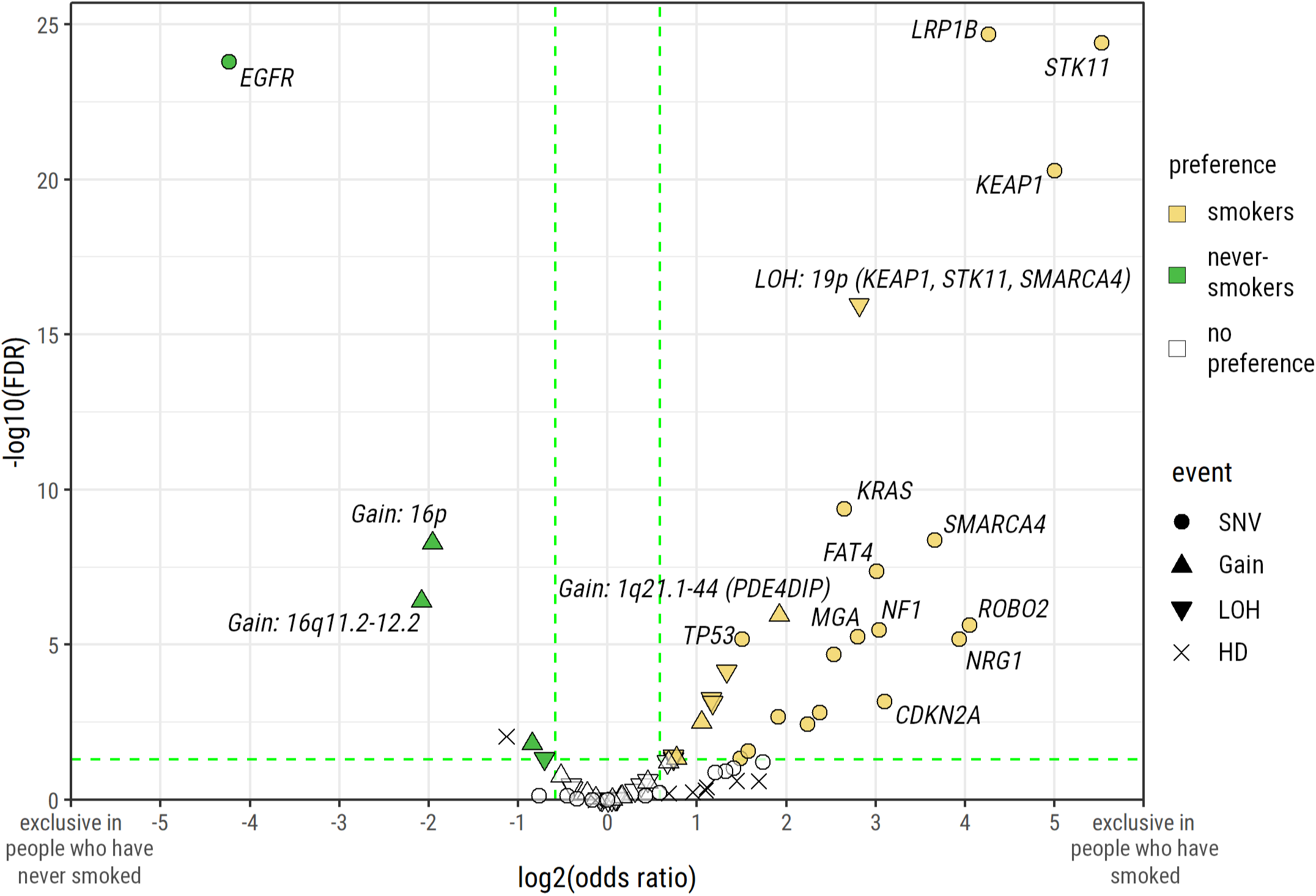
Events showing a significant preference between S-LUAD and NS-LUAD. A volcano plot showing the log2(odds ratio) and -log10(FDR) of event occurrence comparing tumours in people who self-reported as having smoked with those in people who self-reported as having never smoked. Events are coloured if odds ratio > 3/2 or < 2/3, and FDR < 0.05. Events associated with people who have smoked are coloured yellow, and events associated with people who have not smoked are coloured green. SNVs are represented by circles, copy number gains by upward pointed triangles, losses by downward pointed arrows, and HDs by crosses. Due to the number of events meeting the positive association criteria, events are only labelled if odds ratio > 8 or < 1/8, or if FDR < 1^-5^.

**Figure S5:**
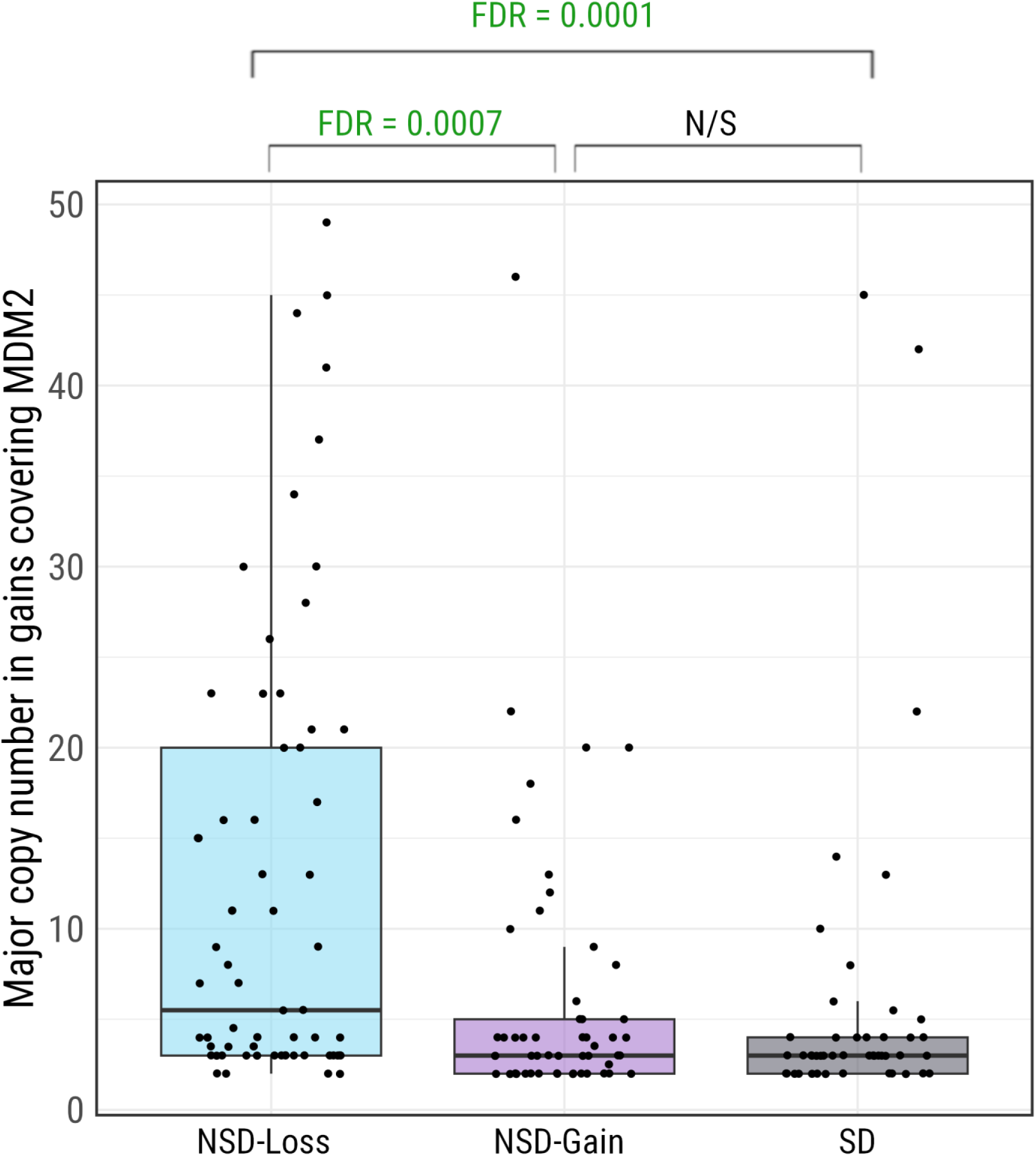
Major copy number of gains covering the MDM2 locus across different trajectories. Boxplots show the major copy number in segments identified as gains, covering the locus of the MDM2 gene. Major copy number is shown for tumours on the NSD-Loss trajectory (blue, left), NSD-Gain trajectory (purple, centre), and SD trajectory (grey, right). FDRs shown relate to multiple testing corrected Wilcoxon tests between each pair of trajectories. Significant FDRs are shown in green.

**Figure S6:**
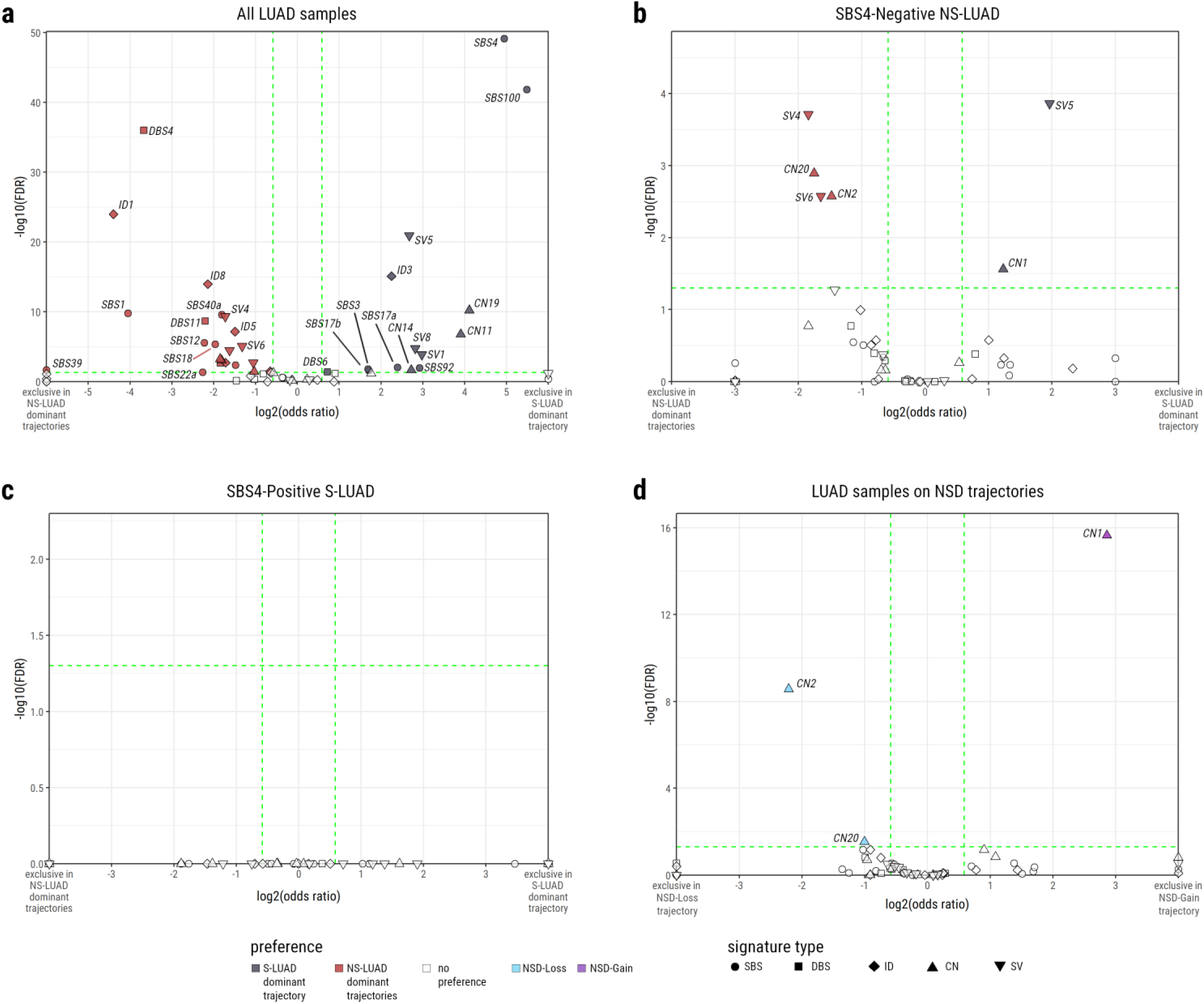
Mutational signatures showing a significant preference between trajectories. Volcano plots show the log2(odds ratio) and -log10(FDR) of single base substitution (SBS), double base substitution (DBS), indel (ID), copy number (CN), and structural variant (SV) signature activity comparing: a) All LUAD between the SD ordering and the NSD-Loss & NSD-Gain orderings combined. b) NS-LUAD between the SD ordering and the NSD-Loss & NSD-Gain orderings combined. c) S-LUAD between the SD ordering and the NSD-Loss & NSD-Gain orderings combined. d) All LUAD between the NSD-Loss ordering and the NSD-Gain ordering. Signatures are coloured if odds ratio > 3/2 or < 2/3, and FDR < 0.05. Signatures are coloured according to their positive associations as follows: SD trajectory (vs NSD trajectories combined) - black, NSD trajectories combined (vs SD trajectory) - red, NSD-Loss (vs NSD-Gain) - blue, NSD-Gain (vs NSD-Loss) - purple, no association - white. SBS signatures are represented by circles, DBS signatures are represented by squares, ID signatures are represented by diamonds, CN signatures are represented by upward pointing triangles, and SV signatures are represented by downward pointing triangles.

**Figure S7:**
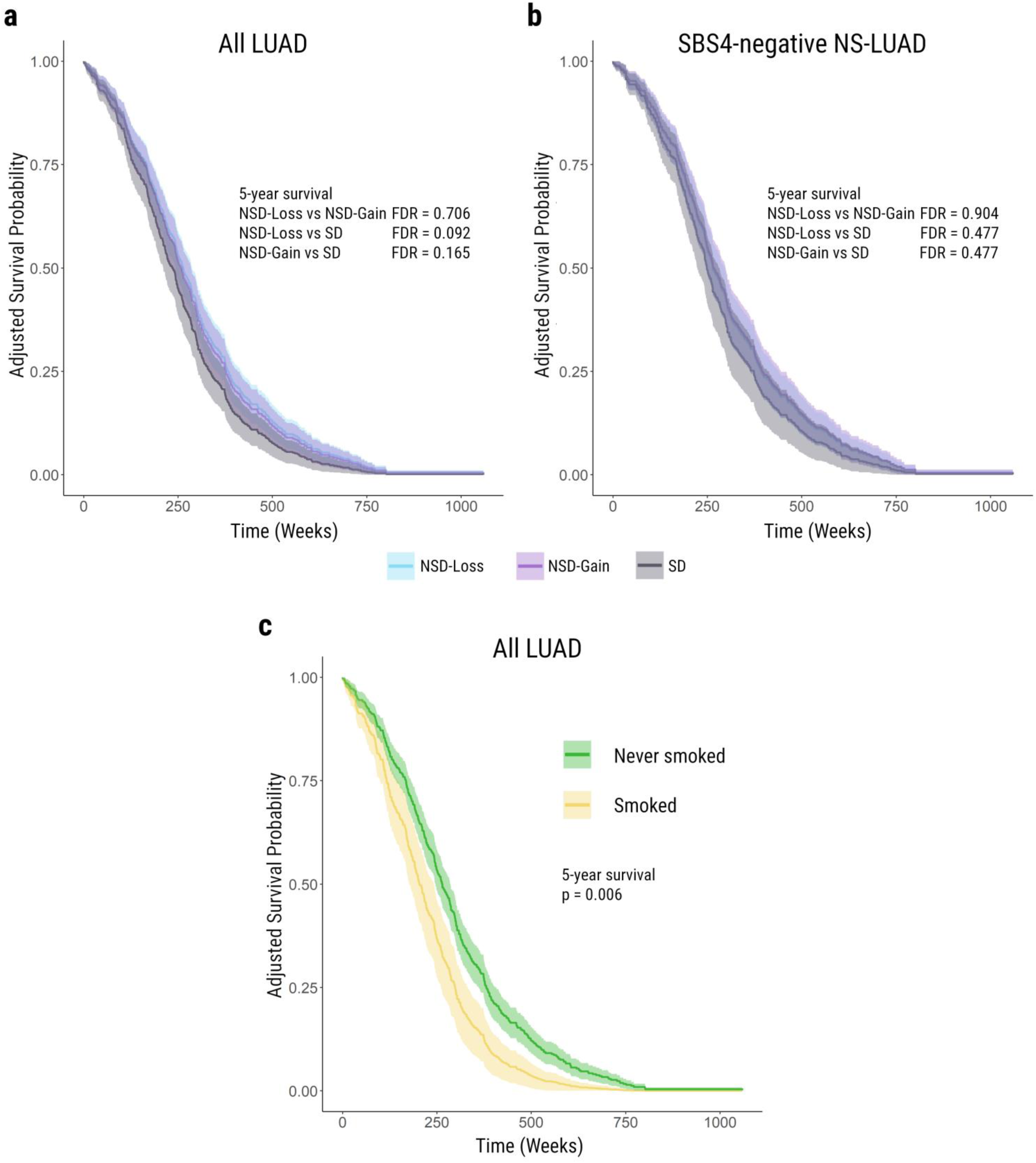
Adjusted survival curves by trajectory and smoking status. Survival curves adjusted for age, sex, and tumour stage. a) shows survival of individuals of all smoking statuses, comparing between NSD-Loss, NSD-Gain, and SD trajectories. b) shows survival of NS-LUAD subjects, comparing between NSD-Loss, NSD-Gain, and SD trajectories. c) shows survival of all individuals, comparing people who self-reported as having smoked with people who self-reported as having never smoked. Significance tests are adjusted curve tests - modified Pepe and Flemming tests for the difference between two adjusted survival curves - at 5 years. NSD-Loss curves are shown in blue, NSD-Gain curves are purple, SD curves are grey, the curve for those who have smoked is yellow, and the curve for those who have never smoked is green.

**Figure S8:**
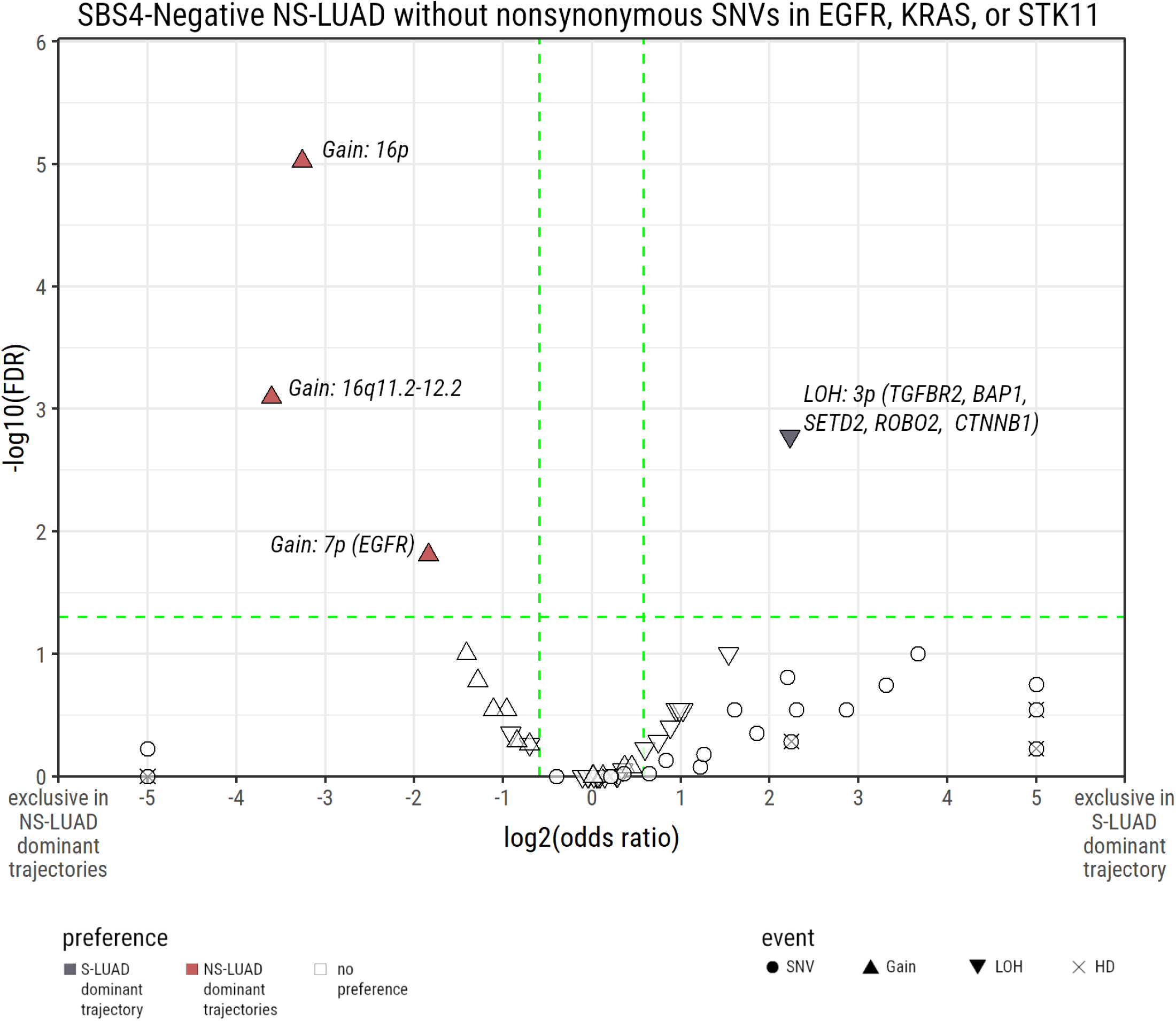
NS-LUAD event preferences in the absence of *EGFR*, *KRAS*, or *STK11* mutations. A volcano plot showing the log2(odds ratio) and -log10(FDR) of event occurrence comparing NS-LUAD between the *SD* ordering and the *NSD-Loss* & *NSD-Gain* orderings combined. Events are coloured if odds ratio > 3/2 or < 2/3, and FDR < 0.05. Events associated with the SD trajectory are coloured black and those associated with the NSD trajectories combined are coloured red. SNVs are represented by circles, copy number gains by upward pointed triangles, losses by downward pointed arrows, and HDs by crosses.

**Figure S9:**
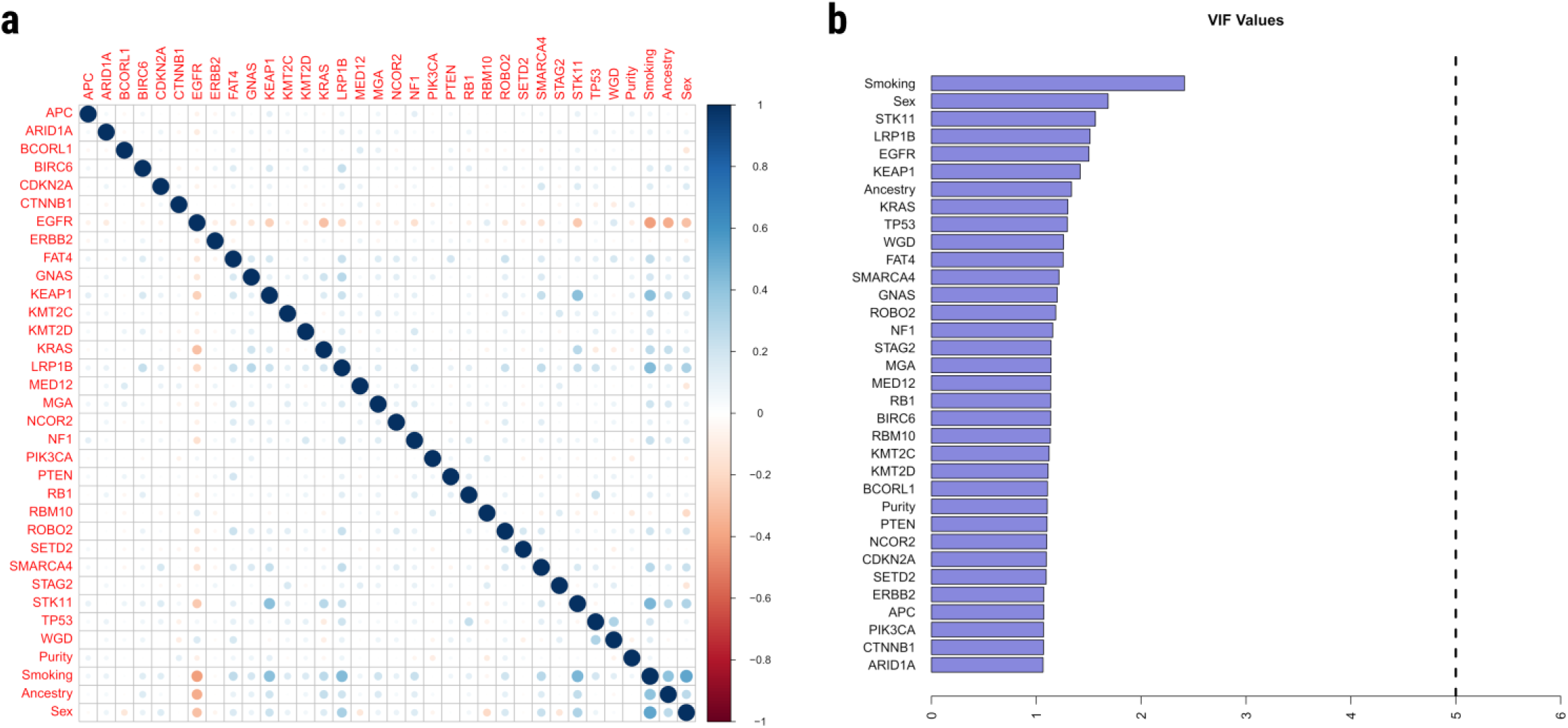
Multicollinearity tests. a) correlation between each pair of variables and b) variance inflation factor (VIF) of each variable to be included in multivariate linear analysis, in order to check that these variables were not subject to too high a degree of multicollinearity. In (a), positive correlations are shown in blue and negative correlations are shown in red. Larger correlations are represented both by darker colours and by larger circles. No pair of variables have a correlation score greater than 0.6. In (b), barplots show VIF values of each variable. A dashed line shows a threshold of 5. No variable has a VIF value greater than 3.

## Author Contributions

Conceptualization: DCW, TZ, MTL; Methodology: CW, MD-G, CDS, YY, LY, BZ, LBA, DCW, MTL.; Formal analysis: CW, YY, XZ, WZ, PHH, JSang, AK, JPM, CH, OWL, SS, KMJ, BZ, MD-G, CDS, TZ, LY, MAN, JShi; Pathology work: CL, MKB, WDT, LMS, PJ, RH, S-RY; Resources: CAH, MPW, KCL, ESE, JMS, MBS, SSY, MManczuk, JL, BS, AM, OS, DZ, IH, DM, MK, YB, BEGR, DCC, VG, PB, GL, PH, NR, DC, QL, SJC, VJ, SO, C-YC, ACP, MSavic, MSaha, I-SC, CW, MTL; Data curation: PHH, TZ, MMiraftab, TVTT, OWL; Writing (original draft): CW, DCW, MTL; Writing (review and editing): CW, PHH, MD-G, LY, MBS, BEGR, DCW, MTL; Visualization: CW; Supervision: DCW, MTL

## Ethics Declarations

Because the National Cancer Institute only received de-identified samples and data from collaborating centres, had no direct contact or interaction with the study participants and did not use or generate identifiable private information, Sherlock-Lung has been determined to constitute ‘not-human-subject research’ (NHSR), on the basis of the federal Common Rule (45 CFR 46; https://www.ecfr.gov/cgi-bin/ECFR?page=browse).

## Acknowledgements

This research was supported by the Intramural Research Program of the National Institute of Health (NIH) (project ZIACP101231 to MTL), and by the Anne Wojcicki Foundation (Grant Number: LC009). The contributions of the NIH author(s) were made as part of their official duties as NIH federal employees, are in compliance with agency policy requirements, and are considered Works of the United States Government. However, the findings and conclusions presented in this paper are those of the author(s) and do not necessarily reflect the views of the NIH or the U.S. Department of Health and Human Services. We want to acknowledge the patients and the INCLIVA Biobank (PT17/0015/0049) integrated in the Spanish National Biobanks Network and in the Valencian Biobanking Network for their collaboration. This study was supported by the Health and Medical Research Fund of Hong Kong SAR, HMRF 03142856. The related studies of Taiwan site were supported by grants from the Ministry of Health and Welfare, Taiwan DOH97-TD-G-111-026 (C.A.H.), DOH98-TD-G-111-015 (C.A.H.), DOH99-TD-G-111-028 (C.A.H.); DOH97-TD-G-111-029 (C.Y.C.), DOH98-TD-G-111-018 (C.Y.C.), DOH99-TD-G-111-015 (C.Y.C.), DOH97-TD-G-111-028(I.S.C.), DOH98-TD-G-111-017(I.S.C.), DOH99-TD-G-111-014(I.S.C.), and the Ministry of Science and Technology, Taiwan MOST109-2740-B-400-002 (C.A.H.), MOST110-2740-B-400-002 (C.A.H.), MOST111-2740-B-400-002 (C.A.H.). This work has been supported in part by the Tissue Core at the H. Lee Moffitt Cancer Center & Research Institute, a comprehensive cancer center designated by the National Cancer Institute and funded in part by a Moffitt Cancer Center Support Grant (no. P30-CA076292). And, in part, by NIH (NCI) grant # U01CA209414 to the Boston Lung Cancer Survival Study of the Dana-Farber/ Harvard Cancer Center (D.C.C.). The authors would like to thank the team at the IUCPQ site of the Quebec Respiratory Health Network Biobank of the FRQS for their valuable assistance, and would like to thank the staff at Harvard University, Yale University, Roswell Park Cancer Institute and Roswell PI, Instituto Nacional de Cancerologia, Nice University Hospital Centre (Nice UHC) - University Côte d’Azur and the Nice Biobank CRB, Toronto University Health Network, and Mayo Clinic for their assistance providing samples and corresponding clinical data. We thank the study participants and the staff at Westat Inc. for their assistance in collecting samples and corresponding clinical data. This work used the computational resources of the NIH HPC Biowulf cluster (http://hpc.nih.gov). DCW is supported by the NIHR Manchester Biomedical Research Centre (NIHR203308). The views expressed are those of the authors and not necessarily those of the NIHR or the Department of Health and Social Care.

## Data availability

Normal and tumour-paired CRAM files for the WGS data for the individuals in the Sherlock-Lung study and the EAGLE study have been deposited in dbGaP under the accession numbers [phs001697.v2.p1](https://www.ncbi.nlm.nih.gov/projects/gap/cgi-bin/study.cgi?study_id=phs001697.v1.p1) and phs002992.v1.p1, respectively. Detailed access information for the publicly available datasets is available in Table S18. Human reference genome GRCh38 was downloaded from the GATK resources at https://github.com/broadinstitute/gatk/blob/master/src/test/resources/large/Homo_sapiens_assembly38.fasta.gz.

## Code availability

Code used for this study can be accessed at https://gitfront.io/r/chriswirth/XC6Q8bvVWfsq/SherlockLung-EvolutionaryTrajectory-Analysis/

